# Regulation of macroautophagy and microautophagic lipophagy by phosphatidylserine synthase Cho1 and external ethanolamine

**DOI:** 10.1101/2025.08.08.669440

**Authors:** Nanaru Mineoka, Rikako Konishi, Yuri Nakashima, Moe Muramoto, Kayoko Fukuda, Sayuri Kuriyama, Tatsunori Masatani, Akikazu Fujita

## Abstract

Phospholipids play crucial roles in autophagy; however, the underlying mechanisms remain elusive. We previously found that the phosphatidylserine (PtdSer) transporter Osh5 is critical for autophagosome formation. Therefore, in this study, we aimed to investigate the impact of the knockout of *cho1*, which encodes PtdSer synthase, on autophagy. Green fluorescent protein-autophagy-related gene 8 (GFP-Atg8) processing assay revealed a significant defect in the macroautophagic activity of the *cho1*Δ mutant, regardless of the presence or absence of ethanolamine (Etn). Notably, autophagosomes were absent in the cytosol, and macroautophagic bodies were not observed in the vacuoles of the starved *cho1*Δ mutant, underscoring the essential role of PtdSer synthesized using Cho1 in autophagosome biogenesis. In contrast, numerous microautophagic vesicles containing lipid droplets were observed in the vacuoles of *cho1*Δ mutants starved in the presence of Etn, suggesting the crucial role of phosphatidylethanolamine (PtdEtn) synthesized via the Kennedy pathway in microautophagic lipophagy when PtdSer synthesis using Cho1 is disrupted. Given recent evidence pointing to the involvement of the ubiquitination system in various autophagy-related processes, we also examined the role of ubiquitin-conjugating enzyme E2 gene *ubc4*. Moreover, microautophagic lipophagy was significantly diminished in starved *cho1*Δ yeast with *ubc4* knockout. These findings suggest the critical role of Ubc4-mediated ubiquitination in facilitating microautophagic lipophagy of the vacuole in yeast cells.

## 1. Introduction

Autophagy involves various systems that transport cytoplasmic components into lysosomes (vacuoles) and perform many physiological and pathological functions [1]. The three main types of autophagy distinguished by their morphological and mechanistic characteristics are macroautophagy, microautophagy, and chaperone-mediated autophagy [1]. Each type involves the delivery of cargo to the lysosomes (vacuoles), where the acidic environment and acid hydrolases break down their contents into basic components for subsequent anabolic processes. Macroautophagy, the most extensively studied form of autophagy, involves the formation of double-membrane vesicles, called autophagosomes, which sequester cellular components. More than 30 autophagy-related genes (ATGs) regulate different stages of autophagy, such as nucleation, orientation, and elongation of the autophagosome and its fusion with the lysosome (vacuole) [2]. In contrast, mechanisms of microautophagy, involving direct invagination of the lysosomal (vacuolar) membrane to engulf the target cargo [3–5], remain unclear.

Lipids are crucial to sustain life, as they are the key components of biological membranes and metabolic energy reserves, contributing significantly to various signaling pathways. Lipids, particularly phospholipids, are also crucial primary membrane sources for the phagophore or isolation membrane and involved in membrane elongation during autophagosome formation in macroautophagy [6–8] and in the expansion of the vacuole membrane during microautophagic lipophagy [9, 10]. In macroautophagy, phospholipid transport by Atg2 [6, 11] and its scrambling by Atg9 [7, 8] are crucial for autophagosome biogenesis. Furthermore, inner membrane (in mammals) or macroautophagic body membrane (in yeast) of the double-membrane structure of the autophagosome undergoes selective degradation after fusion with the lysosome or vacuole [12]. Therefore, phospholipid components are modulated during autophagosome maturation before and during fusion with the target organelles [13–15]. However, the specific types of phospholipids localized or required at each stage of autophagy remain unknown.

Phospholipid distribution in the plasma and intracellular membranes has mainly been studied using fluorescently tagged protein domains binding to hydrophilic head groups of phospholipids in the cytoplasm [16–18]. However, overexpression of these phospholipid-binding protein domains interferes with the membrane recruitment of authentic phospholipid effectors and inhibits phospholipid-dependent cellular processes [19, 20]. For example, overexpression of phosphatidylinositol 3-phosphate (PtdIns(3)P)-binding probes, such as the 2xFYVE domain of EEA1 or PX domain of p40phox, to visualize PtdIns(3)P blocks phagosome maturation at an early stage [21], thus interfering with the analysis of the PtdIns(3)P content of late phagosomes or phagolysosomes. Additionally, this method does not provide information on phospholipid distribution in the luminal leaflets of organelle membranes. Electron microscopy (EM) using ultrathin sections generally detects some lipids in both leaflets [22–24], but it cannot determine the exact location of the lipids in the luminal or cytoplasmic leaflet owing to section thickness and low membrane contrast. Therefore, we previously used the quick-freeze and freeze-fracture replica labeling (QF-FRL) method, which facilitates nanoscale membrane lipid labeling by physically fixing the membrane molecules [13, 25–27]. This technique is particularly important to analyze autophagosomes, in which the double-membranes are closely apposed, and vesicle structures, such as macroautophagic bodies, within the vacuole lumen.

We previously investigated the roles of Osh proteins homologous to oxysterol-binding proteins, which mediate non-vesicular lipid transport, in macroautophagy in yeast [15]. We found that macroautophagic activity was significantly impaired in the *osh1*-*osh7*Δ: *osh4*^ts^ mutant. Additionally, using the QF-FRL method, we found that autophagosomes were absent from the cytoplasm of the mutant. Among the *osh* gene-deletion mutants, *osh5*Δ mutant exhibited a defect in macroautophagic activity, with a notable absence of cytoplasmic autophagosomes. Furthermore, autophagosomes in the *osh5*Δ mutant were smaller, with noticeably thinner inner and outer membranes, compared to those in wild-type (WT) yeast. Labeling density of phosphatidylserine (PtdSer) in the autophagosome membrane of the *osh5*Δ mutant was also significantly lower than that in the autophagosome membrane of WT yeast [15]. These findings indicated the crucial role of PtdSer in autophagosome biogenesis. PtdSer is synthesized using the Cho1 enzyme in yeast cells. In this study, we aimed to further examine the specific roles of PtdSer synthesized using Cho1 in macroautophagy.

Here, our study revealed that PtdSer synthesized using Cho1 was essential for macroautophagy and autophagosome biogenesis in yeast cells. Additionally, exogenously supplied ethanolamine (Etn) induced microautophagic lipophagy in the vacuole of the starved *cho1*Δ mutant, indicating that phosphatidylethanolamine (PtdEtn) synthesized via the Kennedy pathway is essential for the microautophagic lipophagy.

Furthermore, our results suggest the essential role of the ubiquitin-conjugating E2 enzyme Ubc4 in microautophagic lipophagy.

## 2. Methods

### 2.1. Probes

Complementary DNA (cDNA) generated by RT-PCR of HEK293 cell total RNA was used as a template to clone the PX domain of p40Phox and PH domains of OSBP. The plasmid containing the PHx2 domain of Evectin-2 was kindly provided by Dr. Tomohiko Taguchi (Touhoku University) and Dr. Takuma Kishimoto (Hokkaido University). Recombinant GST fusion proteins containing, the PX domain of p40Phox (GST-p40Phox-PX), OSBP PH domain (GST-OSBP-PH), Evectin-2 PH domain (GST-Evectin-2-PH), and phospholipase C (PLC)-δ1 PH domain (GST-PLCδ1-PH) were expressed in *Escherichia coli* and purified as previously described [14, 27, 28]. The PH domain of PLC-δ1 was obtained from the GST-PLC-PH domain fusion protein via cleavage with PreScission protease (GE Healthcare Life Sciences, Pittsburgh, PA, USA) according to the manufacturer’s protocol. The secondary rabbit anti-GST antibody was purchased from Bethyl Laboratories (Montgomery, TX, USA). 10 nm gold particle-conjugated anti-mouse IgG and anti-rabbit IgG antibodies were purchased from BioCell (Cardiff, UK) and Jackson ImmunoResearch (West Grove, PA, USA). Biotin-conjugated duramycin-LC (biotin-duramycin), and mouse anti-biotin antibody were purchased from Polysciences (Warrington, PA, USA), and Jackson ImmunoResearch Laboratories (West Grove, PA, USA), respectively. Rabbit monoclonal anti-GFP antibody (Cell Signaling Technology, Danvers, U.S.A.) and HRP-linked anti-rabbit IgG antibody (Cytiva, Tokyo, Japan) were used for western blotting analysis.

### 2.2. Cultivation of yeast cells

Yeast cells were grown in YPD medium (1% yeast extract, 2% Bacto Peptone and 2% dextrose) or SMD (synthetic minimal medium; 0.67% yeast nitrogen base without amino acids, 2% glucose) at 30 °C, and were harvested during the transition from exponential or stationary phase. All yeast strains used in this study were based on the parent strain *Saccharomyces cerevisiae* SEY6210 (*MATAα leu2-3, 112 ura3-52 his3-Δ200 trp1-Δ901 lys2-801 suc2Δ9 GAL*) [29] (Table S1). Deletion mutants were generated using a polymerase chain reaction (PCR)-based method with pUG6, pAG32, and pFA6a-TRP1 plasmids as templates [30]. To induce autophagy, cells cultured in YPD medium were washed with water and incubated for 3–5 h at 30 °C in nitrogen and carbon-deleted medium S(-NC) (0.17% yeast nitrogen base without amino acids and ammonium sulfate (Bacton Dickinson), nitrogen-depleted medium SD(-N) (0.17% yeast nitrogen base without amino acids and ammonium sulfate and 2% dextrose) or carbon-depleted medium SN(-C) (1% yeast extract and 2% Bacto Peptone). S(-NC), SD(-N) and SN(-C) mediums were supplemented with 1 mM phenylmethylsulfonyl fluoride (PMSF, Sigma) when needed.

### 2.3. Plasmids

The plasmid pRS316-GFP-ATG8 [31] was provided by the National BioResource Project, Japan.

### 2.4. Macroautophagic activity assay

Macroautophagic activity was assessed by monitoring the degradation of GFP-tagged Atg8 within vacuoles, following the method described by Cheong et al. [32].

### 2.5. Quick-freezing and freeze**–**fracture electron microscopy

Quantitative lipid labeling of the biological membranes of budding yeast cells was performed by quick-freezing yeast cells using either a metal sandwich method or high-pressure freezing using an HPM 010 high-pressure freezing machine (Leica Microsystems, Wetzlar, Germany). For metal sandwich freezing, a small volume of yeast pellet was placed on a copper foil, covered with a thin gold foil (∼4 mm^2^ in area; 20 μm thick), and frozen using a quick press between two gold-plated copper blocks precooled in liquid nitrogen [28, 33]. For high-pressure freezing, an EM grid (200 mesh) impregnated with yeast cells was sandwiched between a flat aluminum disc (Engineering Office M. Wohlwend, Sennwald, Switzerland) and frozen according to the manufacturer’s instructions [34].

For freeze–fracture, frozen specimens were transferred to a cold stage of a Balzers BAF400 apparatus (Bal-Tec AG, Lichtenstein) and fractured at −130 °C under a vacuum of ∼1 × 10^−6^ millibars. Replicas were prepared by electron-beam evaporation in three steps: carbon (C) (∼2-nm-thick) at an angle of 90° to the specimen surface, platinum–carbon (Pt/C; 1–2 nm) at an angle of 45°, and C (10–20 nm) at an angle of 90°, as previously described [25]. The thickness of the deposit was adjusted using a crystal-thickness monitor (EM QSG100, Leica).

Thawed specimens were incubated with 2.5% sodium dodecyl sulfate (SDS) in 0.1 M Tris–HCl (pH 8.0) at 60 °C to 70 °C overnight. To remove the cell wall, yeast replicas were digested with 1 mg/mL Zymolyase 20T in phosphate-buffered saline (PBS) containing 0.1% Triton X-100, 1% bovine serum albumin (BSA), and a protease inhibitor cocktail (Nacalai Tesque) for 2 h at 37 °C. After further treatment with 2.5% SDS, the replicas were stored in buffered 50% glycerol at –30 °C until use.

### 2.6. Labeling and electron microscopy observation

Probe labeling was performed as described previously [25]. Briefly, after rinsing, freeze-fracture replicas were blocked with PBS containing 2% cold-water fish skin gelatin at room temperature for 30 min. To label PtdIns(3)P, PtdIns(4)P, and PtdSer, the samples were incubated at 4 °C overnight with 30 μg/mL GST-p40Phox-PX, 40 μg/mL GST-OSBP-PH, or 20 μg/mL GST-Evectin2-PH in PBS containing 2% cold-water fish skin gelatin, respectively. Then, the samples were successively incubated in PBS containing 2% cold-water fish skin gelatin at 37 °C for 30 min each with rabbit anti-GST antibody (5 μg/mL) and 50-fold diluted colloidal gold (10 nm)-conjugated anti-rabbit IgG antibody (EM.GFAR 10; BBI Solutions, Cardiff, UK). To block the nonspecific binding of GST-OSBP-PH to PtdIns(4,5)P_2_, the replicas were pretreated with the PH domain (1 mg/mL) of PLC-δ1 in a blocking solution [28]. To label PtdEtn, biotin-duramycin (60 μM) was diluted in PBS containing 2% cold-water fish skin gelatin. After four washes with PBS containing 0.1% BSA, replicas were incubated at 37 °C for 30 min with mouse anti-biotin antibody (130 μg/ml), followed by incubation with 10 nm gold-conjugated anti-mouse IgG antibody (Jackson ImmunoResearch, West Grove, PA, USA) in PBS containing 2% cold-water fish skin gelatin.

### 2.7. Statistical Analysis

Each experiment was repeated at least three times. The areas on the images were measured using Fiji (ImageJ), and the labeling density was indicated as the number of colloidal gold particles in 1 μm^2^. For each structure, the labeling density was measured in more than 10 randomly selected images. Statistical differences between samples were tested using *t*-test or Dunnett’s multiple tests.

## 3. Results

### 3.1. Defects in macroautophagic activity and autophagosome formation in the cho1 deletion mutant

To observe macroautophagic membranes, yeast cells were cultured for 5 hours in S(-NC) medium supplemented with PMSF, which is devoid of nitrogen and carbon sources. This condition was chosen for subsequent experiments as it robustly induces macroautophagy, facilitating frequent observation of larger autophagosomes [35]. In agreement with previous studies [34, 36], freeze-fracture EM revealed autophagosomes in yeast as spherical structures surrounded by double smooth membranes (Fig. 1A) [37]. The outer and inner membranes of autophagosomes were clearly distinguishable based on the convexity/concavity of their structure and their relative positions within the double membrane (Fig. 1A). Smooth-surfaced macroautophagic bodies were also identified within the vacuolar lumen (blue and green in Fig. 1B), consistent with the notion that these structures are derived from the inner autophagosomal membrane. In autophagy-deficient mutants lacking genes such as ATG2, ATG3, ATG5, ATG7, ATG8, ATG9, ATG12, ATG16, ATG18, VPS15, or VPS34, neither autophagosomes nor macroautophagic bodies were observed as we previously reported [36]. Similar results were obtained when macroautophagy was induced by nitrogen starvation using SD(-N) medium [36] or carbon starvation using SD(-Dextrose) medium (unpublished data, [38]) in the presence of PMSF. Moreover, macroautophagic bodies were largely absent from the vacuoles of yeast cells starved in S(-NC) medium without PMSF, consistent with prior observations [15]. In contrast, they accumulated markedly in the vacuoles of *pep4*Δ and *atg15*Δ mutants under the same conditions, even in the absence of PMSF, as previously reported [14]. These findings suggest that macroautophagic bodies are normally degraded by vacuolar proteinases and lipases. We previously reported that the vacuoles of starved WT yeast predominantly contain two types of vesicles̶macroautophagic bodies and microautophagic vesicles̶arising from macroautophagy and microautophagic lipophagy, respectively [9, 10, 14]. However, distinguishing between them based on morphological features is challenging, as both vesicle types exhibit smooth surfaces and similar sizes when observed using the freeze-fracture EM method (see Figs. 1B, 2A). As reported previously [14], our quick-freezing and freeze-fracture replica labeling (QF-FRL) method revealed that PtdIns(3)P was localized in both the PF (cytoplasmic leaflet, see Supplemental Fig. S1) and EF (luminal leaflet) of yeast autophagosomal membranes (Fig. 1A). Labelling of PtdIns(3)P was significantly stronger in the EF of both the inner and outer membranes. Additionally, PtdIns(3)P was intensively labeled in the membranes of macroautophagic bodies, showing a similar luminal leaflet (EF)-dominant asymmetry (Figs. 1B, and S2). These findings support the hypothesis that macroautophagic bodies are derived from the inner autophagosomal membrane (Supplementary Fig. S3), which predominantly features PtdIns(3)P in its EF (Supplementary Fig. S4). In contrast to macroautophagic bodies, we and other groups previously reported that PtdIns(3)P is localized in the PF, but not EF, in microautophagic vesicles in the vacuole caused by microautophagic lipophagy (see Figs. 2 and S2) [10, 14].

**Figure 1.**
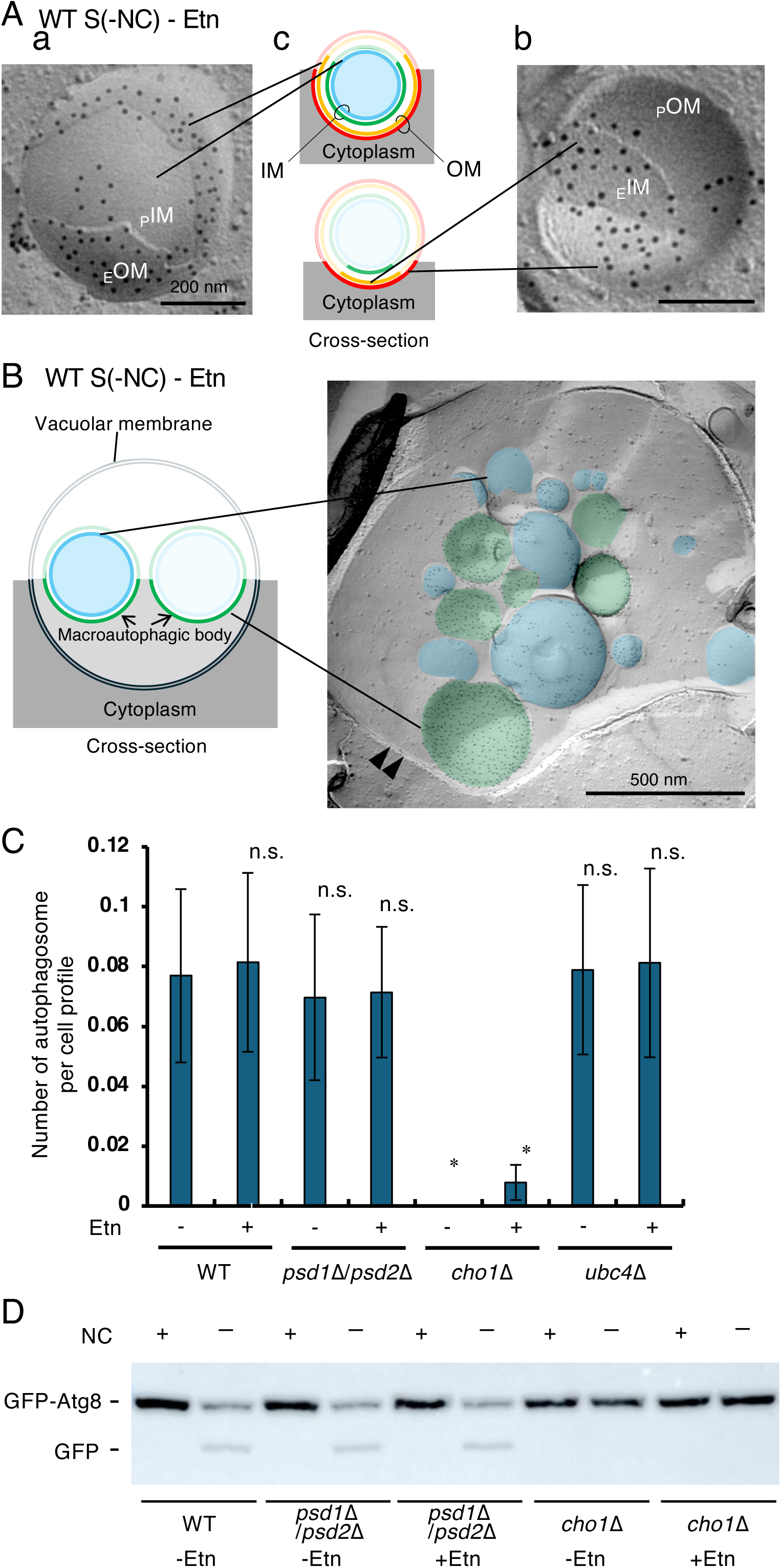
Defects in autophagosome formation and autophagic activity in *cho1*Δ yeast. (A) Autophagosomes in wild-type (WT) cultured in S(-NC) for 5 hr. The autophagosome was identified as s smooth-surfaced double-membrane structure, and seen in two different patterns, as shown in the diagrams in (c). (a) Convex structure displaying the outer membrane (OM) EF (yellow line, _E_OM) and the inner membrane (IM) PF (blue line, _P_IM). (b) Concave structure displaying the OM PF (red line, _P_OM) and the IM EF (green line, _E_IM). The labelings of PtdIns(3)P were detected in the EF of both the outer (a) and inner (b) autophagosome membranes. Scale bar, 200 nm. (B) WT yeast cultured in S(-NC) supplemented with PMSF for 5 h. PtdIns(3)P was labelled intensely in the EF (green) of macroautophagic body membranes in the vacuolar lumen. A lower amount of labelling was also seen in the PF (blue) of macroautophagic body membranes. Double arrowheads, vacuole. Scale bar, 500 nm. (C) The freeze-fracture EM showed that autophagosomes were observed in the cytosol of WT, *psd1*Δ/*psd2*Δ, and *ubc4*Δ strains after incubation in S(-NC) for 5 h. However, almost no autophagosomes were detected in the cytosol of *cho1*Δ mutant strain in both the presence and absence of Etn in the S(-NC) medium. Mean ± standard error (SE) of three independent experiments (>35 cells were counted in each group in each experiment). The number of autophagosomes per cell in each strain was compared to that in WT yeast. Dunnett’s test: n.s., not significant; \**p* < 0.05. (D) Green fluorescent protein (GFP)-autophagy-related gene (Atg)-8 degradation via macroautophagy in the *cho1*Δ mutant. Wild-type (WT), *psd1*Δ/*psd2*Δ, and *cho1*Δ strains transformed with a plasmid expressing GFP-Atg8 under the control of the *CUP1* promoter were grown in SMD-Ura medium at 30 °C in the presence or absence of 3 mM ethanolamine (Etn) to mid-log phase. For each strain, the culture was divided into three aliquots. One aliquot was cultured in YPD medium for 4 h at 30 °C. The other aliquots were shifted to S(-NC) and cultured for 4 h at 30 °C in the presence or absence of Etn. For each aliquot, samples were taken before (+NC) and after (-NC) starvation without PMSF. Bands of full-length GFP-Atg8 and free GFP are shown. Defect in GFP-Atg8 degradation was observed in the *cho1*Δ mutant in both the presence and absence of Etn.

**Figure 2.**
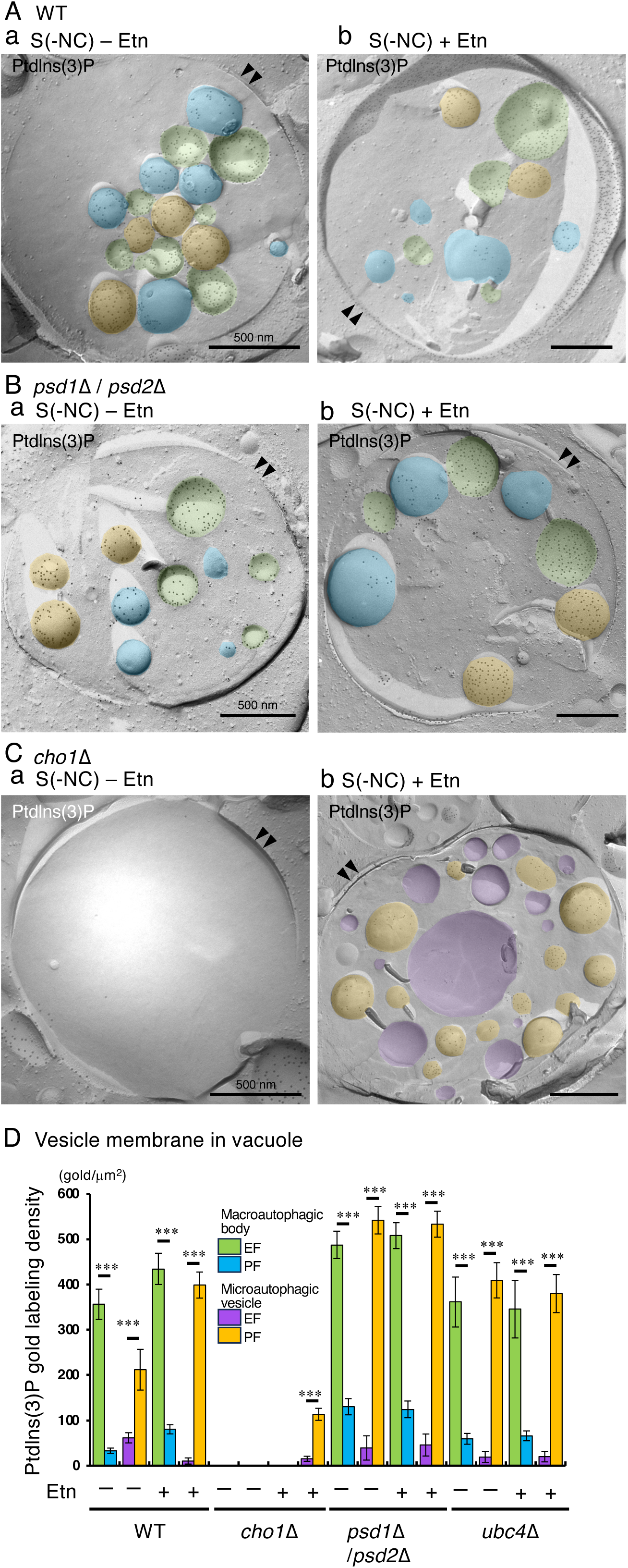
Phosphatidylinositol 3-phosphate (PtdIns(3)P) labeling of vesicle membranes in the vacuoles of starved WT yeast, *psd1*Δ/*psd2*Δ, *cho1*Δ, and *ubc4*Δ mutants. WT yeast (A), *psd1*Δ/*psd2*Δ (B), *cho1*Δ (C), and *ubc4*Δ mutant strains were starved in S(-NC) medium supplemented with PMSF for 5 h at 30 °C, both in the absence (a) and presence (b) of ethanolamine (Etn). In WT yeast (A), and the *psd1*Δ/*psd2*Δ (B) and *ubc4*Δ mutants, two types of vesicles were observed under both conditions: one with PtdIns(3)P labeling enriched in the external face (EF, green), but not cytoplasmic face (PF, blue), of vesicles in the vacuoles and another with labeling in the peripheral face (PF, orange), but not the EF (purple) (D, see also Supplementary Fig. S2). In contrast, no vesicles were detected in the vacuoles of the *cho1*Δ mutant in the absence of Etn (Ca). However, in the presence of Etn, numerous vesicles were observed in the vacuoles of the *cho1*Δ mutant (Cb). Unlike WT yeast, and the *psd1*Δ/*psd2*Δ and *ubc4*Δ mutants, only one type of vesicle was detected in the *cho1*Δ mutant with Etn supplementation. In these vesicles, PtdIns(3)P labeling was localized to the PF (orange) but not the EF (purple) of the vesicle membranes in the vacuoles (Cb, D). This characterization of the PtdIns(3)P labelings indicates that these vesicles in the vacuole of the *cho1*Δ mutant were microautophagic vesicles. The microautophagic vesicles were classified when the labeling densities of PtdIns(3)P were more than 200 golds/μm^2^ in the PF and less than 50 golds/μm^2^ in the EF of vesicular membrane. Mean ± SE of three independent experiments (>10 vacuoles were counted in each group in each experiment). PtdIns(3)P labeling in the PF was compared with that in the EF. *t-*test: *** *p* < 0.001. Scale bar: 500 nm.

The ULK complex (Atg1 in yeast) and FIP200 (Atg11 and Atg17 in yeast) play a central role in autophagy initiation by assembling on the ER membrane, a critical hub for phospholipid synthesis required for autophagosome formation [39]. Notably, FIP200 has been observed to colocalize with key enzymes involved in phospholipid biosynthesis, such as phosphatidylserine synthase 1 (PSS1) and phosphatidylinositol synthase (PIS). PIS synthesizes phosphatidylinositol (PI), which serves as a precursor for PtdIns(3)P [40] and PtdIns(4)P [36, 41], both of which are indispensable for macroautophagy. Additionally, our previous findings demonstrated that the macroautophagic activity was significantly impaired in the *osh1*-*osh7*Δ: *osh4*^ts^ mutant, which lacks functional PtdSer transporters. The cytoplasmic abundance of autophagosomes was markedly reduced in the *osh5*Δ mutant, and PtdSer labeling in autophagosome membranes was substantially lower compared to WT yeast, if detectable at all [15]. These results suggest that PtdSer itself or its metabolic derivatives, synthesized by Cho1, likely play a pivotal role in the regulation of macroautophagy in budding yeast.

To explore the role of Cho1, the PtdSer synthase, in macroautophagy, we analyzed the macroautophagic membranes of *cho1Δ* mutants using freeze-fracture EM. We found in this study that the appearance rate of autophagosomes in the cytoplasm was significantly lower in starved *cho1Δ* mutants, regardless of the presence or absence of Etn, compared to the WT strain (Fig. 1C). Moreover, macroautophagic bodies were absent in the vacuoles of the starved *cho1*Δ mutants (Figs. 2C and 3D). Atg8, a ubiquitin-like protein crucial for autophagosome formation, relies on its conjugation to PtdEtn for association with autophagosomal membranes [42, 43]. In yeast, PtdEtn is primarily synthesized through two pathways: from PtdSer via Psd1 [44, 45] or Psd2 [46], and via the cytidine diphosphate (CDP)-Etn (Kennedy) pathway [47] (Supplementary Fig. S5). Additionally, minor pathways contribute to PtdEtn synthesis in yeast [48]. In this study, freeze-fracture EM showed that autophagosomes were present in the cytoplasm of starved WT yeast cells, both with and without Etn (Fig. 1C). Similarly, the marked difference in autophagosome formation between *cho1*Δ and *psd1*Δ/*psd2*Δmutants indicates that the presence of PtdSer itself, rather than its conversion to downstream metabolites, is crucial for autophagosome formation and macroautophagy (Fig. 1C). Macroautophagic bodies were also detected in the vacuoles of both starved WT and *psd1*Δ*/psd2*Δ strains in S(-NC) medium containing PMSF, regardless of Etn supplementation (Figs. 2 and 3D). These observations indicate that PtdEtn synthesized via both the major and minor pathways is sufficient for Atg8 conjugation. However, the results highlight that PtdSer, synthesized by the Cho1 enzyme, is indispensable for autophagosome formation.

**Figure 3.**
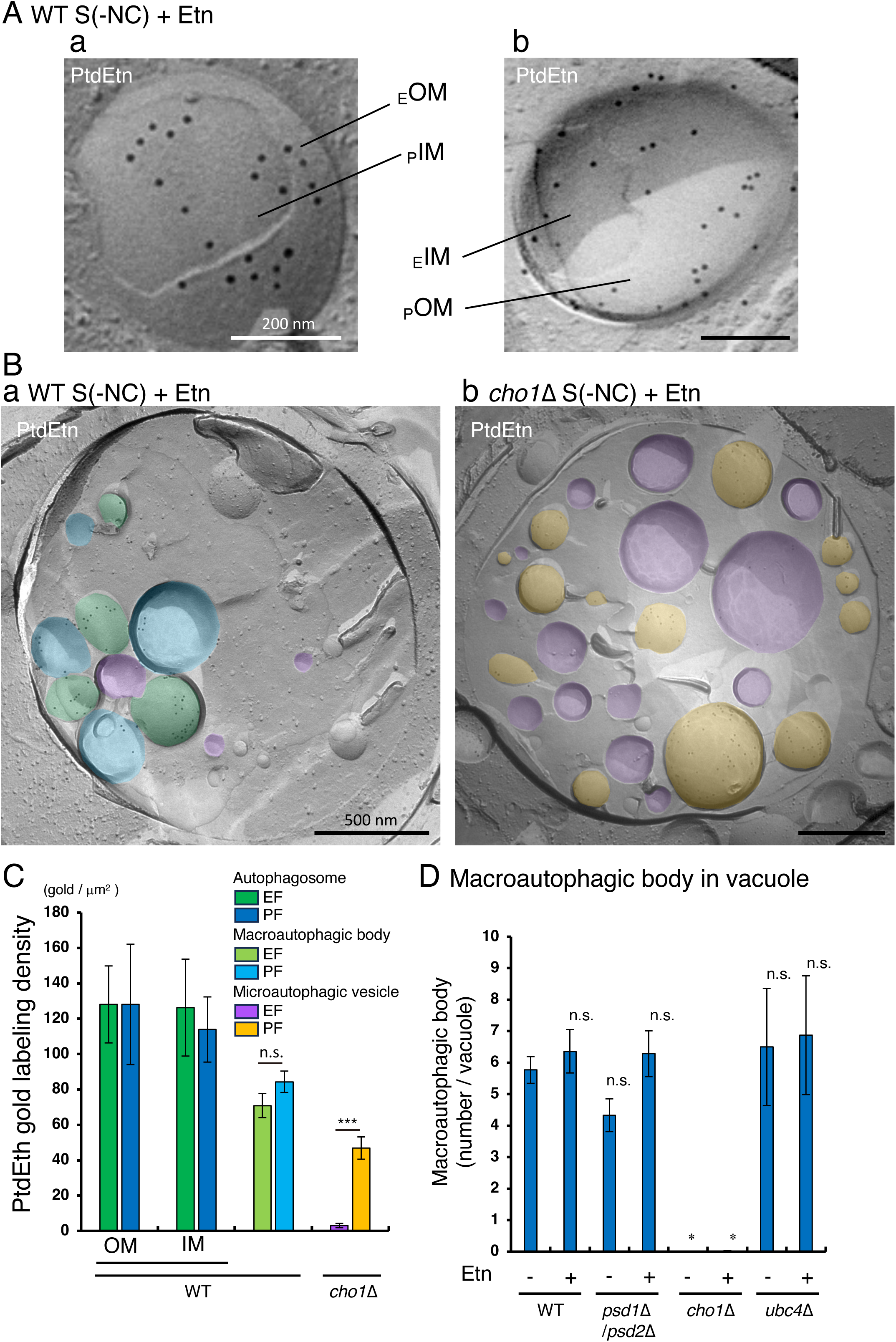
PtdEtn labeling of autophagosome membrane, macroautophagic body, and microautophagic vesicle in WT yeast, the *psd1*Δ/*psd2*Δ, *cho1*Δ, and *ubc4*Δ mutants. WT, *psd1*Δ/*psd2*Δ, *cho1*Δ, and *ubc4*Δ yeast strains were starved in the S(-NC) medium supplemented with PMSF in the absence and presence of Etn at 30 °C for 5 h. Microphotographs of autophagosomes (A), macroautophagic bodies, and microautophagic vesicles in the vacuoles of WT yeast (Ba) and the *cho1*Δ mutant (Bb) are shown. PtdEtn labeling was detected in both the EF and PF of the inner and outer membranes of autophagosomes (A) in the cytoplasm and macroautophagic bodies (Ba) in vacuoles. In the *cho1*Δ mutant, PtdEtn labeling was localized in the PF, but not the EF, in the membrane of the microautophagic vesicles in the vacuoles (Bb). Gold labeling densities of EF were not significantly different from those of PF in the membranes of macroautophagic bodies (C). IM: inner membrane, OM: outer membrane. Mean ± SE of three independent experiments (>30 autophagosomes or vesicles were counted in each group in each experiment). PtdEtn labeling in PF was compared to that in EF. *t-*test; n.s., not significant; \*\*\**p* < 0.001. (D) According to the localization pattern of PtdIns(3)P (Fig. 2) and PtdEtn labeling in EF or PF of the vesicular membranes, the numbers of the macroautophagic bodies in the vacuoles of WT, *psd1*Δ/*psd2*Δ, *cho1*Δ, and *ubc4*Δ strains were determined. Several macroautophagic bodies were detected in WT yeast and *psd1*Δ/*psd2*Δ in the presence and absence of the Etn, but almost none were detected in the vacuoles of *cho1*Δ mutant in both the presence and absence of Etn. Mean ± SE of three independent experiments (>20 vacuoles were counted in each group in each experiment). The number of macroautophagic bodies in the vacuoles of each strain was compared to that in WT yeast in the absence of Etn. Dunnett’s test: n.s., not significant; \**p* < 0.05.

Atg8, a ubiquitin-like protein, is conjugated to PtdEtn and remains associated with mature autophagosomes as described above. During macroautophagy, green fluorescent protein (GFP)-tagged Atg8 is degraded in the vacuoles, leading to the release of free GFP. The presence of a relatively stable GFP component serves as an indicator of macroautophagy. We confirmed the deficiencies of the macroautophagy in the *cho1*Δ mutant using this GFP-Atg8 processing assay. Here, we did not detect free GFP under nutrient-rich conditions. However, after cultured the cells in the S(-NC) medium without PMSF for 4 h, a distinct band of free GFP was visible in WT yeast (Fig. 1D). In this study, normal macroautophagic activity was observed in WT yeast cells starved in S(-NC) medium in the absence of Etn (Fig. 1D). Furthermore, *psd1*Δ/*psd2*Δ mutant exhibited comparable macroautophagic activity to WT yeast in both the presence and absence of Etn (Fig. 1D). These results align with a previous report [49], which found no detectable defects in the macroautophagic activity of starved *psd1*Δ and *psd2*Δ mutants, suggesting that alternative PtdEtn synthesis pathways can compensate for the loss of specific pathways. In contrast, a significantly lower amount of free GFP was observed in the *cho1*Δ strain than in the WT strain (Fig. 1D), indicating a significant deficiency in macroautophagy in the *cho1*Δ mutant in both the presence and absence of Etn (Fig. 1D). Similar results were obtained in all strains starved by nitrogen starvation using SD(-N) medium for 4 h (Supplementary Fig. S6). These results are consistent with the observation obtained with the freeze-fracture EM method.

### 3.2. Microautophagic lipophagy in the starved cho1Δ mutant

In the starved *cho1*Δ mutant, no vesicular structures were detected in the vacuole lumen using the freeze-fracture EM method when starvation was induced in S(-NC) containing PMSF in the absence of Etn (Fig. 2Ca). However, many vesicular structures were observed in the vacuole lumen of the starved *cho1*Δ mutant in the presence of Etn (Fig. 2Cb). As described above, the QF-FRL method revealed that PtdIns(3)P is predominantly localized in EF of the macroautophagic body (Fig. 1B), which originates from the inner vesicle of the autophagosome, in the vacuole (Supplementary Fig. S3). In this study, we detected PtdIns(3)P labeling in the PF, but not in EF, of the vesicular membranes within the vacuole lumen (Fig. 2Cb) of the *cho1*Δ mutant starved in the presence of Etn, indicating that these vesicular structures are not identical to the macroautophagic bodies. Moreover, several macroautophagic bodies, in which the PtdIns(3)P labeling was predominantly detected in EF, but not in PF, were detected in the vacuoles of WT yeast and *psd1*Δ/*psd2*Δ mutant starved in the absence and presence of Etn (Fig. 2A, B, and D). In addition to PtdIns(3)P, PtdEtn labeling was detected in both the EF and PF of the autophagosomes (Fig. 3A) and macroautophagic bodies (Fig. 3Ba) of starved WT yeast. Labeling densities of PtdEtn were not significantly different between the EF and PF of the autophagosome in the cytoplasm (Fig. 3A, C) and macroautophagic bodies in the vacuole (Fig. 3Ba, C) of starved WT yeast. However, PtdEtn labeling was localized in PF, but not in EF, of the vesicles in the vacuolar lumen of the *cho1*Δ mutant starved in the presence of Etn (Fig. 3Bb and C).

Labeling density of PtdEtn in PF was significantly higher than that in EF of the vesicles in the vacuolar lumen of the starved *cho1*Δ mutant in the presence of Etn (Fig. 3Bb, C). Therefore, macroautophagic body was not observed in the vacuolar lumen of the starved *cho1*Δ mutant both in the absence and presence of Etn (Figs. 2C and 3D).

Freeze-fracture EM has revealed that the vacuolar membrane exhibits characteristic geometric patterns during the stationary phase [9, 10, 50], and that domains lacking the transmembrane protein Vph1 correspond to raft-like regions [10, 51, 52]. In the present study, freeze-fracture EM demonstrated that the vacuolar membrane of the *cho1Δ* mutant exhibited such geometric patterns when starved in the presence, but not in the absence, of Etn (Fig. 4A; Supplemental Fig. S6). Notably, invaginated, intramembrane particle (IMP)-deficient raft-like domains, tightly associated with lipid droplets (LDs; indicated by white double arrowheads in Fig. 4B), were observed under these conditions. Furthermore, the IMP-rich, non-raft-like regions were confined to the periphery of these raft-like protrusions, forming a distinct ring-like structure at the neck of the balloon-shaped domain in the vacuolar membrane (arrows in Fig. S8; [10]).

**Figure 4.**
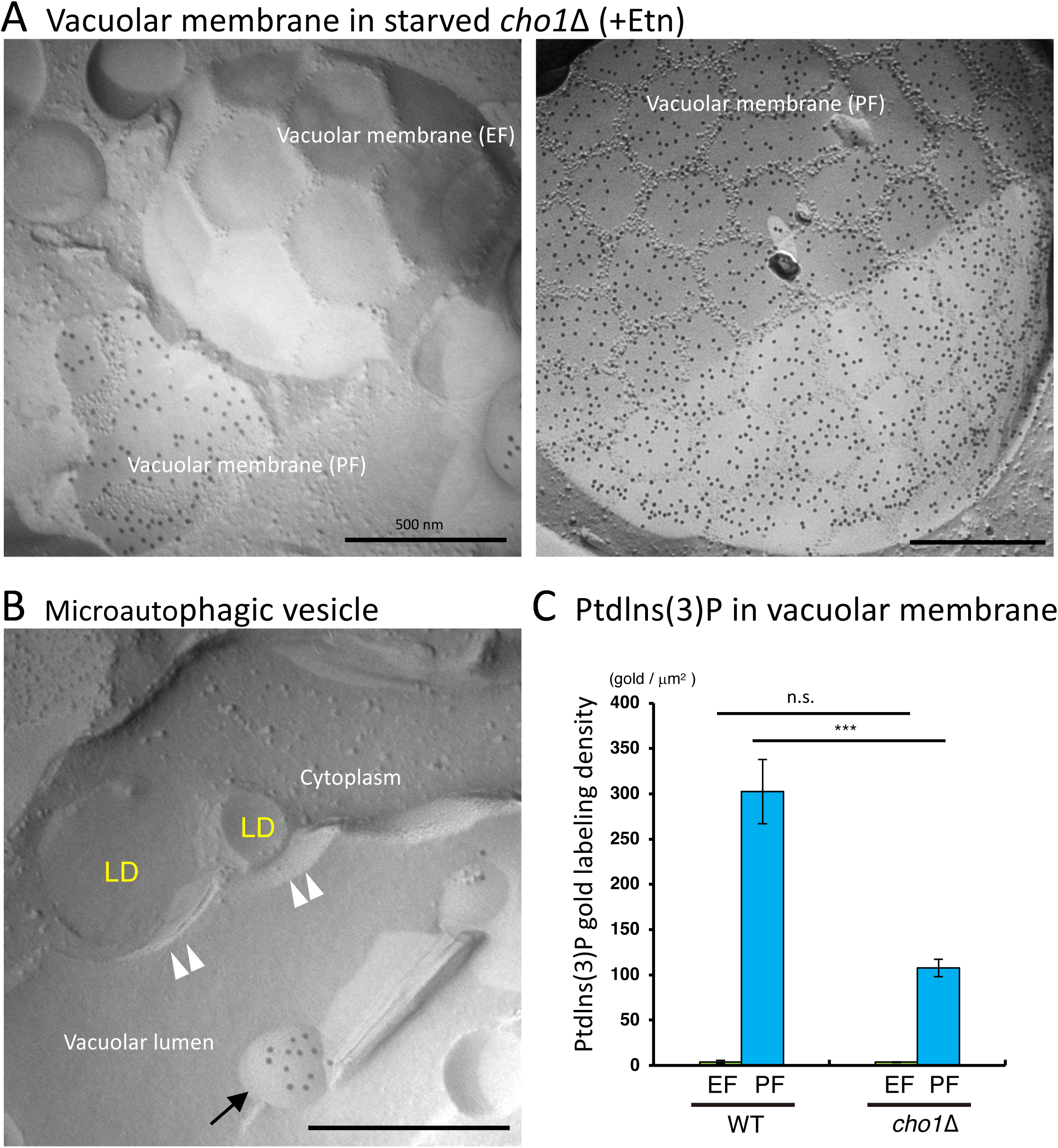
Microautophagic lipophagy in the starved *cho1*Δ mutant occurs via the expansion of raft-like vacuolar membrane in the presence of Etn. *cho1*Δ mutant yeast was incubated in S(-NC) medium supplemented with PMSF in the presence of Etn at 30 °C for 5 h. Freeze-fracture EM method and PtdIns(3)P labeling of the vacuolar membrane (A and B). Vacuolar membrane in *cho1*Δ mutant starved in the presence of the Etn showed geometrical patterns in both EF (left panel in A) and PF (right panel in A). Intra-membrane particles (IMPs), which are transmembrane proteins, were largely confined to the edges of polygonal areas. (B) Lipid droplets (LDs) adhered tightly to the IMP-deficient raft-like domain, which bulged toward the vacuolar lumen. The vacuolar membrane and microautophagic vesicles are indicated by double arrowheads and arrows, respectively. Scale bar: 500 nm. (C) Labeling densities of PtdIns(3)P in the vacuolar membrane of WT yeast and *cho1*Δ mutant. PtdIns(3)P labeling was abundant in PF, and its density in the *cho1*Δ mutant was much lower compared to that in WT yeast (see also Supplementary Fig. S10). Mean ± SE of three independent experiments (>20 vacuoles were counted in each group in each experiment). PtdIns(3)P labeling in EF and PF was compared to that in WT yeast. *t-*test: n.s., not significant; \*\*\**p* < 0.001.

Clusters of IMPs were also identified within vesicles in the vacuolar lumen (arrows in Fig. S8), suggesting that these raft-like protrusions are severed at the non-raft regions, completing the process of microautophagic lipophagy (Supplementary Fig. S3).

Previous studies have shown that PtdIns(3)P [10] and PtdIns(4)P [9] localize specifically to the PF, but not the EF, of microautophagic vesicles in the vacuole. In agreement with this, we found that in *cho1Δ* mutants starved in the presence of Etn, both PtdIns(3)P (Fig. 2D) and PtdIns(4)P (Supplementary Fig. S9) were predominantly localized to the PF of vesicles within the vacuole. Notably, the labeling density of PtdIns(3)P on the PF of these vesicles was markedly reduced in *cho1Δ* cells compared to WT [10, 14] and *psd1Δ/psd2Δ* mutants (Fig. 2A, B, D; Supplementary Fig. S2). A similar reduction in PtdIns(3)P labeling was observed in the vacuolar membrane itself: while PtdIns(3)P was consistently localized to the PF in WT cells [34], its density was significantly lower in *cho1Δ* mutants (Fig. 4C; Supplementary Fig. S10). These findings indicate that the vesicles present in the vacuolar lumen originate from the vacuolar membrane and are microautophagic in nature (Supplementary Figs. S3, S8, and S11).

Consistently, using the QF-FRL method, we detected several microautophagic vesicles in the vacuolar lumen of *cho1Δ* mutants starved in the presence, but not the absence, of Etn (Fig. 5B). Vps27, a core component of the ESCRT-0 complex, is essential for endosomal sorting [53] and has also been implicated in vacuolar microautophagic lipophagy [54, 55]. In our study, no microautophagic vesicles were detected in the vacuoles of *vps27Δ/cho1Δ* mutants, even when starved in the presence of Etn (Fig. 5Ad). Furthermore, the proportion of vacuolar membranes displaying geometric patterns was drastically reduced in *vps27Δ/cho1Δ* mutants under these conditions (Fig. 6B, C). Collectively, these results suggest that microautophagic lipophagy is specifically induced in *cho1Δ* mutants only when ethanolamine is present during starvation.

**Figure 5.**
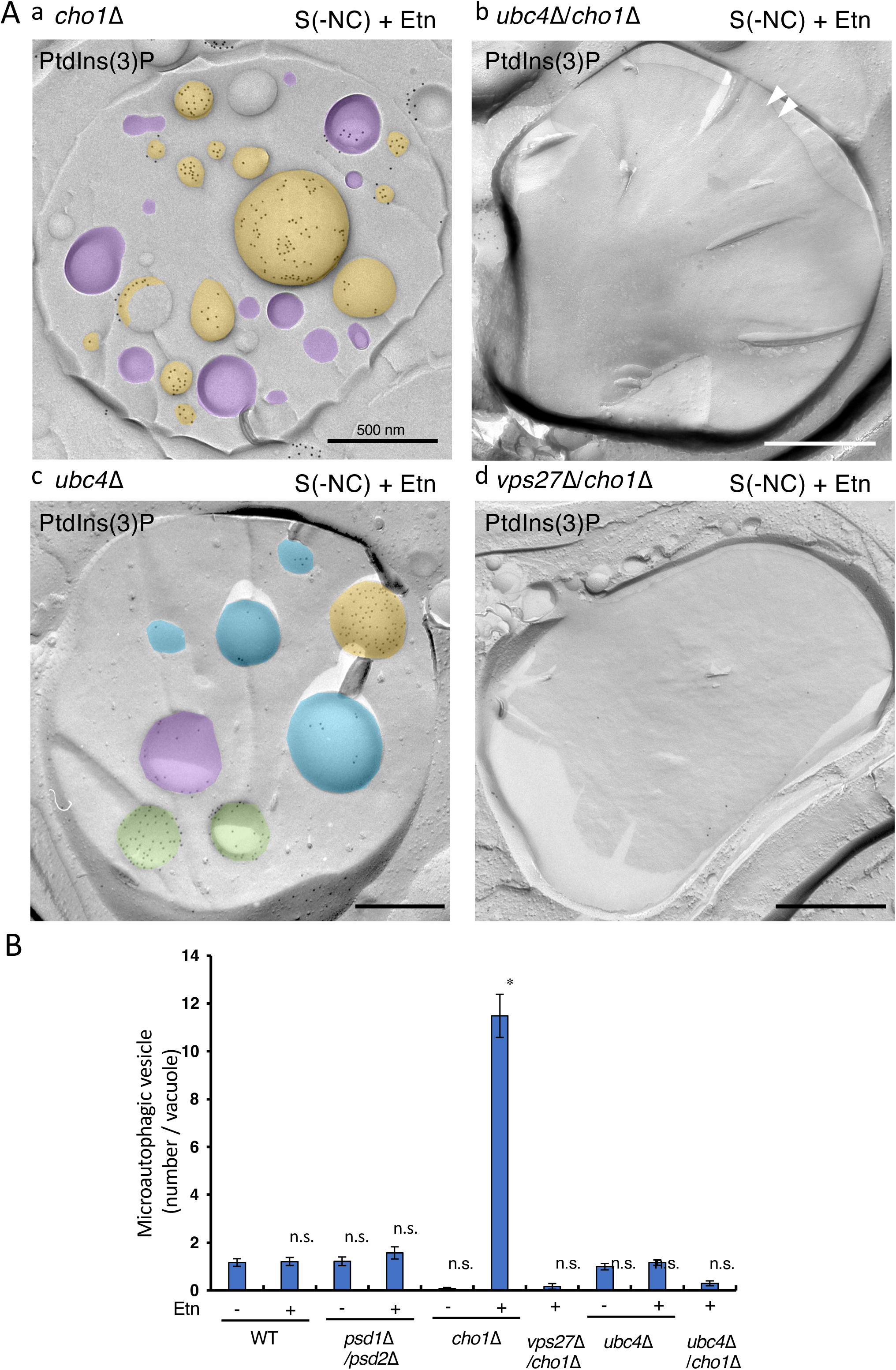
Microautophagic lipophagy in the starved *cho1*Δ mutant depends on the expression of Vps27. (A) Micrographs depicting PtdIns(3)P labeling of macroautophagic bodies (blue and green) and microautophagic vesicles (yellow and purple) within the vacuoles of *cho1*Δ (a), *ubc4*Δ/*cho1*Δ (b), *ubc4*Δ (c), and *vps27*Δ/*cho1*Δ (d) mutants starved supplemented with PMSF in the presence of Etn. Notably, no microautophagic vesicles were detected in the vacuoles of starved *ubc4*Δ/*cho1*Δ (b) and *vps27*Δ/*cho1***Δ** (d) mutants, even with Etn supplementation. (C) According to the localization patterns of PtdEtn (Fig. 3) and PtdIns(3)P (Fig. 2) labeling in EF and PF of the vesicular membranes, the numbers of microautophagic vesicles in the vacuoles of the WT, *psd1*Δ/*psd2*Δ, *cho1*Δ, *vps27*Δ/*cho1*Δ, *ubc4*Δ, and *ubc4*Δ/*cho1*Δ strains were determined. Several microautophagic vesicles were detected in the vacuolar lumen of the starved *cho1*Δ mutant in S(-NC) supplemented with PMSF in the presence (+Etn), but not in the absence (-Etn) of Etn. Mean ± SE of three independent experiments (>30 vacuoles were counted in each group in each experiment). The number of microautophagic vesicles in the vacuole of each strain was compared to that in WT yeast. Dunnett’s test: n.s., not significant; \**p* < 0.05.

**Figure 6.**
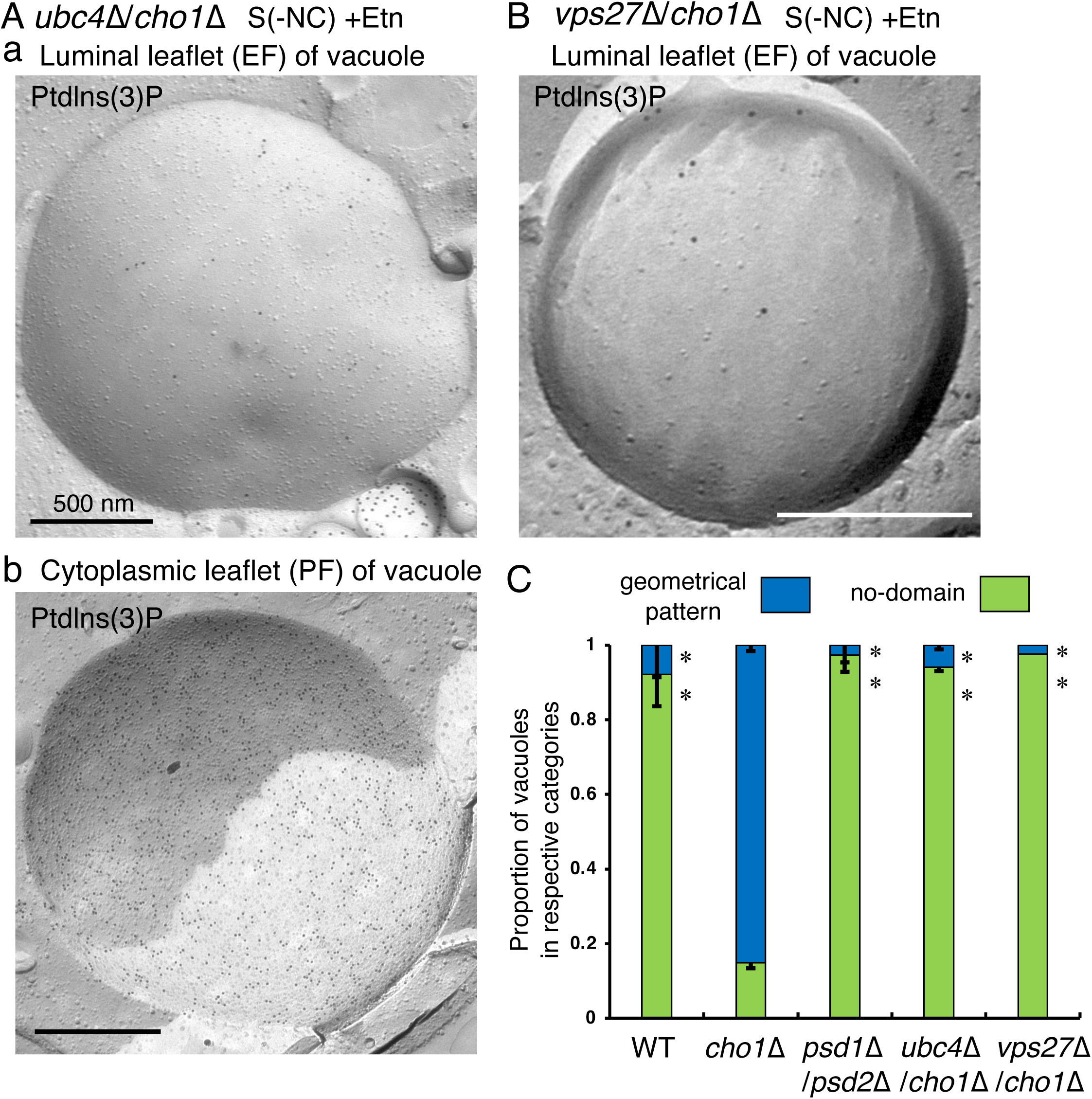
Ubc4 is required for geometrical pattern of the vacuolar membrane. Micrographs of the PtdIns(3)P labeling in the vacuolar membrane of the *ubc4*Δ/*cho1*Δ (A) and *vps27*Δ/*cho1*Δ (B) mutants starved supplemented with PMSF are shown. When the *ubc4*Δ/*cho1*Δ and *vps27*Δ/*cho1*Δ mutants were starved in the presence of Etn, the geometrical pattern was rarely observed in both the luminal (a) and cytoplasmic (b) leaflets of the vacuolar membrane. Scale bar: 500 nm. (C) Proportion of geometrical and no-domain patterns in the vacuolar membranes of WT yeast and the *cho1*Δ, *psd1*Δ/*psd2*Δ, *ubc4*Δ/*cho1*Δ, and *vps27*Δ/*cho1*Δ mutants starved in the presence of Etn. Ratios of the geometrical patterns in the starved *ubc4*Δ/*cho1*Δ and *vps27*Δ/*cho1*Δ mutants were drastically decreased. Mean ± standard deviation (SD) of three independent experiments (>100 vacuoles were counted in each group in each experiment). Ratios of each pattern in WT, *psd1*Δ/*psd2*Δ, *ubc4*Δ/*cho1*Δ, and *vps27*Δ/*cho1*Δ mutants were compared to that in the *cho1*Δ mutant. Dunnett’s test: \**p* < 0.05.

### 3.3. Ubiquitin-conjugating E2 enzyme Ubc4 is required for microautophagic lipophagy

Sakamaki et al. [56] recently reported that ubiquitin is conjugated to phospholipids, mainly PtdEtn, in yeast and mammalian cells and showed that ubiquitinated PtdEtn accumulates in the endosome and vacuole/lysosome membranes. Furthermore, they reported that PtdEtn ubiquitination is facilitated by starvation in yeast cells [56]. In this study, exogenously supplied Etn induced microautophagic lipophagy in the starved *cho1*Δ mutant (Figs. 4 and 5), indicating that PtdEtn ubiquitination plays a role in microautophagic lipophagy in the starved *cho1*Δ mutant. PtdEtn ubiquitination is significantly reduced in the ubiquitin-conjugating enzyme (E2) gene *ubc4*-knocked-out yeast [56]. Geometrical pattern of the vacuolar membrane, which was detected in the *cho1*Δ mutant starved in the presence of Etn, was not observed in the starved *ubc4*Δ/*cho1*Δ mutant in the presence of Etn (Fig. 6A, C). Furthermore, number of microautophagic vesicles was drastically reduced in the vacuole of the *ubc4*Δ/*cho1*Δ mutant starved in the presence of Etn (Fig. 5Ab, B). These findings emphasize the essential role of the ubiquitin-conjugating E2 enzyme Ubc4 in microautophagic lipophagy.

Our group and others [9, 10, 57] have reported that microautophagic lipophagy of the vacuole occurs in acutely starved WT yeast, accompanied by the formation of IMP-deficient raft-like domains in the vacuolar membrane. Microautophagic vesicles were observed in the vacuoles of both WT and *psd1*Δ/*psd2*Δ mutants starved in S(-NC) medium supplemented with PMSF, regardless of the presence or absence of Etn (Fig. 2A, B; Supplementally Fig. S2). Unexpectedly, microautophagic vesicles were also detected in the vacuole of the starved *ubc4*Δ mutant (Fig. 5Ac). Furthermore, in acutely starved WT yeast [10] and *ubc4*Δ mutant (Supplementally Fig. S12), raft-like domains were restricted to specific regions of the vacuolar membrane, whereas in the stationary phase [9, 10] and in the starved *cho1*Δ mutant supplemented with Etn (Fig. 4), the vacuole was almost entirely covered with these domains. These findings suggest that Ubc4 is not essential for vacuolar microautophagic lipophagy under acute starvation conditions in WT yeast cells.

## 4. Discussion

In this study, we mainly found that (i) macroautophagy was defective in the *cho1*Δ mutant, (ii) autophagosome was absent in the starved *cho1*Δ mutant, (iii) microautophagic lipophagy was induced in the *cho1*Δ mutant starved in the presence of Etn, and (iv) microautophagic lipophagy was inhibited in the starved *ubc4*Δ/*cho1*Δ mutant in the presence of Etn.

### 4-1. Essential roles of PtdSer synthesized by Cho1p in the autophagosome formation

In our previous study [15], we demonstrated the significant reduction in the number of autophagosomes in the *osh5*Δ mutant compared to that in the WT strain, suggesting the critical role of Osh5 in autophagosome formation. Additionally, we reported thin inner and outer membranes of the autophagosome, if any, in the *osh5*Δ mutant, along with low PtdSer labeling on the autophagosome membrane, emphasizing the significance of PtdSer localization in autophagosome biogenesis [15]. Based on these findings, we investigated the macroautophagic activity of the *cho1*Δ mutant using the GFP-Atg8 processing assay in this study. We observed a significant defect in the macroautophagic activity of the *cho1*Δ mutant, as evidenced by the absence of autophagosomes in the cytoplasm following starvation, regardless of the presence or absence of Etn (Fig. 1). These findings underscore the essential role of the PtdSer synthase, Cho1, in autophagosome biogenesis.

In yeast cells, PtdEtn is synthesized via main two distinct pathways (Supplementary Fig. S5). In one pathway, PtdEtn is produced from PtdSer by the decarboxylases, Psd1 [44, 45] and Psd2 [46]. In the other pathway, PtdEtn is synthesized via the Kennedy pathway using Etn and diacylglycerol [47, 58]. A previous report suggested that PtdEtn is rarely formed in the *cho1*Δ mutant yeast in the absence of Etn [59], in spite of the existence of other minor synthetic pathways of PtdEtn [48] (Supplementary Fig. S5). Consistently, growth defects have been reported in the *cho1*Δ mutant in the absence of Etn [59, 60]. PtdEtn is crucial for macroautophagy as its conjugation with Atg8 is an essential step in autophagosome formation [42, 61]. In this study, we observed significant macroautophagic activity, and autophagosomes in the cytoplasm and many macroautophagic bodies in the vacuoles of WT yeast even in the absence of Etn and *psd1*Δ/*psd2*Δ mutant in the presence or absence of Etn (Fig. 1).

These results were well consistent with the previous report in which the macroautophagic activities were not inhibited in the *psd1*Δ or *psd2*Δ mutants [49]. These findings suggest that PtdEtn synthesized through both major and minor pathways (Supplementary Fig. S5) contributes to yeast macroautophagy. In contrast to macroautophagy, previous studies have shown that Psd1, but not Psd2, is essential for mitochondria-specific autophagy (mitophagy) in yeast under acute nitrogen starvation [49]. This indicates that PtdEtn, produced from PtdSer by the Psd1 decarboxylase localized in the mitochondrial inner membrane [45], plays a crucial role in mitophagy by facilitating Atg8 conjugation for autophagosome formation. In this study, we found that macroautophagic activity was mostly inhibited, and no autophagosomes in the cytoplasm or macroautophagic bodies in the vacuoles were observed in the starved *cho1*Δ mutant, even in the presence of Etn (Figs. 1 and 3). These results indicate that autophagosome formation does not occur even in the presence of PtdEtn, when PtdSer is not synthesized using Cho1. To the best of our knowledge, this is the first study to show that PtdSer synthesized using Cho1 is essential for autophagosome formation in yeast cells. However, further studies are necessary to elucidate the action mechanisms of PtdSer in autophagosome formation.

### 4-2. Inducible microautophagic lipophagy in the starved cho1Δ mutant

Here, microautophagic lipophagy was induced in the starved *cho1*Δ mutant yeast when Etn was exogenously added to the medium. This indicates the important role of exogenous Etn in microautophagic lipophagy. As described above, two major pathways (Supplementary Fig. S5), one involving the conversion of PtdSer by Psd1 [44, 45] or Psd2 [46] and the other involving the Kennedy pathway [47], are involved in PtdEtn synthesis; however, additional minor pathways may also exist in yeast cells [48]. Yeast cells with mutations in *cho1* gene require exogenous Etn for normal growth [59]. In this study, we added Etn to the medium to culture the *cho1*Δ mutant as its growth was extremely slow in the absence of Etn (see the Methods section), similar to previous reports [59, 60]. *cho1* mutants exhibit low levels of PtdEtn in the absence of Etn but normal levels of PtdEtn in the presence of Etn [59]. Therefore, exogenously supplied Etn facilitates the synthesis of PtdEtn in the *cho1*Δ mutant, and the synthesized PtdEtn is essential for the induction of microautophagic lipophagy under starved conditions.

Additionally, Vps27 is the central factor of the ESCRT-0 complex in yeast cells [53] and we observed almost no microautophagic vesicles in the vacuoles of the *vps27*Δ/*cho1*Δ mutant starved in the presence of Etn (Fig. 5Ad). These results are consistent with a previous report [54, 55] that microautophagic lipophagy, which is induced after a diauxic shift or a starvation, is abolished in the *vps27* gene-deletion mutant. Here, microautophagic lipophagy was induced in the starved *cho1*Δ mutant in the presence, but not in absence, of Etn. To the best of our knowledge, this is the first study to show that PtdEtn, synthesized via the Kennedy pathway, plays a crucial role in microautophagic lipophagy.

### 4-3. Crucial roles of ubiquitin-conjugating E2 enzyme Ubc4 for microautophagic lipophagy

Sakamaki et al. [56] used a biochemical technique and reported that ubiquitinated PtdEtn is localized in the endosome-or vacuole/lysosome-rich membrane fractions, suggesting its localization in the membranes of endosomes and vacuoles/lysosomes. They also showed PtdEtn ubiquitination in the endosomal and vacuolar membranes is facilitated by starvation [56]. In yeast cells, PtdEtn ubiquitination is catalyzed by the ubiquitin-activating enzyme (E1), Uba1, ubiquitin-conjugating enzymes (E2s), Ubc4 and Ubc5, and ubiquitin ligase (E3), Tul1 [56]. In this study, we found that the microautophagic lipophagy was defective in the starved *ubc4*Δ/*cho1*Δ mutant in the presence of Etn (Figs. 5 and 6). PtdEtn ubiquitination is strongly suppressed in the *ubc4*Δ mutant [56]. These results suggest that PtdEtn ubiquitination plays a crucial role in microautophagic lipophagy in yeast cells.

Liposomes containing ubiquitinated PtdEtn specifically recruit the ESCRT components, Vps27-Hse-1 (ESCRT-0) and Vps23 (a component of ESCRT-1), in vitro [56].

Consistently, in this study, no microautophagic vesicles were observed in the vacuoles of the starved *vps27*Δ/*cho1*Δ mutant in the presence of Etn (Fig. 5Ad).

In this study, PtdEtn was localized in the PF, but not the EF, leaflets of the vacuolar membrane, which exhibited a geometrical pattern in the starved *cho1*Δ mutant in the presence of Etn (Supplementary Fig. S13). Conversely, PtdEtn was localized to both the PF and EF of the vacuolar membrane and exhibited a normal round pattern in starved WT yeast. PtdEtn labeling density in EF, but not in PF, of the vacuolar membrane with the geometric pattern in the starved *cho1*Δ mutant was significantly lower than that in starved WT yeast (Supplementary Fig. S13B). Although whether the PtdEtn-specific binding probe, duramycin, binds to ubiquitinated PtdEtn remains unclear, PtdEtn localized in the cytoplasmic leaflet of the vacuolar membrane is readily ubiquitinated. The ubiquitinated PtdEtn localized in the cytoplasmic leaflet further recruits the ESCRT components, Vps27-Hse1 and Vps23 [56]. Here, vacuolar membranes with geometric patterns were mostly absent in the *ubc4*Δ/*cho1*Δ and *vps27*Δ/*cho1*Δ mutants starved in the presence of Etn (Fig. 6). Formation of an IMP-deficient raft-like domain in the vacuolar membrane with geometric pattern is crucial for the invagination of LDs into the vacuolar lumen, facilitating microautophagic lipophagy [9, 10] (Supplementary Fig. S8). These findings suggest that the binding of Vps27 to ubiquitinated PtdEtn is essential for the formation of an IMP-deficient raft-like domain in the vacuolar membrane exhibiting a geometric pattern. Ubiquitinated PtdEtn on vacuoles is recognized by various ESCRT components, including Vps23 and Hse1, ultimately leading to the recruitment of downstream ESCRT components and inward membrane invagination [62]. However, the ubiquitin-conjugating E2 enzyme Ubc4 can ubiquitinate not only PtdEtn but also various other substrates [63]. Indeed, Vps27, which contains ubiquitin-interacting motifs (UIMs), has been reported to bind ubiquitinated proteins at the endosome [64, 65]. Furthermore, we did not directly assess the ubiquitination level of PtdEtn under our experimental conditions. Therefore, we cannot rule out the possibility that Ubc4-mediated ubiquitination of molecules other than PtdEtn contributes to the microautophagic lipophagy of vacuoles in the starved *cho1*Δ mutant in the presence of Etn. Further investigations are required to clarify the precise mechanisms by which ubiquitinated PtdEtn induces microautophagic lipophagy.

In this study, we demonstrated that microautophagic vesicles were present in the vacuole of acutely starved WT yeast and the *psd1*Δ/*psd2*Δ mutant, regardless of the presence of Etn (Fig. 2). These findings align well with previous reports from our group and others [9, 10, 57]. However, while microautophagic lipophagy was observed in the acutely starved *cho1*Δ mutant with Etn, it was inhibited in the *ubc4*-deletion mutant (*ubc4*Δ/*cho1*Δ, Fig. 5), despite remaining intact in the starved *ubc4*Δ mutant (Fig. 5).

Additionally, we observed that IMP-deficient raft-like domains were partially present in the vacuolar membrane of acutely starved WT yeast [9, 10] and *ubc4*Δ mutant (Supplementary Fig. S12). In contrast, these domains entirely covered the vacuolar membrane in the starved *cho1*Δ mutant with Etn (Fig. 4, Supplementary Fig. S8) [9, 10].

These results suggest that the mechanisms underlying microautophagic lipophagy may differ depending on whether microautophagic lipophagy is associated with the geometric pattern of IMP-deficient raft-like domains. Further investigations will be necessary to elucidate the precise mechanisms governing microautophagic lipophagy.

### 4-4. Formation of IMP-deficient raft-like domain in the vacuolar membrane and microautophagic lipophagy

It has been reported that IMP-deficient raft-like domains in the vacuolar membrane play critical roles in microautophagic lipophagy during both stationary phase and acute starvation [9, 10, 52]. Cholesterol transport to the vacuolar membrane via Niemann-Pick type C (NPC) proteins—NPC1 and NPC2 (Ncr1 and Npc2 in yeast, respectively)—is essential for the expansion of these IMP-deficient raft-like domains [10]. In mammalian cells, intraluminal vesicles (ILVs) within multivesicular bodies (MVBs) have been shown to be enriched in sterols [66]. Furthermore, ILVs are frequently observed in the vacuoles of acutely starved *ncr1*Δ/*npc2*Δ mutants and in mutants lacking the major vacuolar proteases Pep4 and Prb1 [10]. Notably, the formation of IMP-deficient raft-like domains and microautophagic lipophagy during stationary phase is absent in *vps4*Δ mutants, in which the MVB pathway is disrupted [10]. In line with these findings, we also observed pronounced accumulation of ILVs and autophagic bodies within the vacuolar lumen of starved *pep4*Δ mutants (Supplemental Fig. S14), whereas such structures were only rarely found in WT yeasts. These observations support the idea that ILVs within MVBs represent a major sterol source for NPC-mediated expansion of IMP-deficient raft-like domains in the yeast vacuolar membrane [10]. In the present study, we further demonstrated that both IMP-deficient raft-like domain formation (Fig. 6) and microautophagic lipophagy (Fig. 5) were largely abolished in starved *vps27*Δ/*cho1*Δ and *ubc4*Δ/*cho1*Δ mutants in the presence of Etn. It is well established that Vps27 (ESCRT-0) is essential for the MVB pathway in endosomes [62], and that PtdEtn ubiquitinated by Ubc4 can recruit Vps27 to the endosomal membrane [56]. We therefore hypothesize that blocking sterol transport from the MVB to the vacuolar membrane leads to the loss of IMP-deficient raft-like domain formation and microautophagic lipophagy in starved *vps27*Δ/*cho1*Δ and *ubc4*Δ/*cho1*Δ mutants, indicating an indirect role for Vps27 and Ubc4 in these processes.

Additionally, it was shown that raft-like domain formation and microautophagic lipophagy were only partially diminished in *are1*Δ/*are2*Δ mutants, which are defective in sterol ester synthesis (see Figure 6-figure supplement 2 in [10]), suggesting that mechanisms other than cholesterol transport also contribute to the expansion of IMP-deficient raft-like domains in the yeast vacuolar membrane. We further propose that ubiquitination of specific molecules by Ubc4 may directly facilitate the expansion of raft-like domains and promote microautophagic lipophagy in the yeast vacuole. Further studies will be necessary to elucidate the molecular mechanisms underlying the expansion of IMP-deficient raft-like domains in the vacuolar membrane and the process of microautophagic lipophagy in yeast cells.

## Abbreviations

BSA: bovine serum albumin
CDP: cytidine diphosphate
DAG: diacylglycerol
EF: E-face (exoplasmic leaflet)
EM: electron microscopy
ESCRT: endosomal sorting complex required for transport
Etn: ethanolamine
GFP: green fluorescent protein
GST: glutathione S-transferase
HRP: horse radish peroxidase
IMP: intra-membrane particle
ORF: open reading frame
Osh: oxysterol-binding protein homologue
PBS: phosphate-buffered saline
PF: P-face (protoplasmic or cytoplasmic leaflet)
PH: pleckstrin homology
PtdEtn: phosphatidylethanolamine
PtdSer: phosphatidylserine
PtdIns(3)P: phosphatidylinositol 3-phosphate
PtdIns(4)P: phosphatidylinositol 4-phosphate
PtdIns(4,5)P_2_: phosphatidylinositol 4,5-bisphosphate
QF: quick-freezing
QF-FRL: quick-freezing and freeze-fracture replica labeling
RT-PCR: reverse transcription-polymerase chain reaction
SDS: sodium dodecyl sulfate
YPD: yeast peptone dextrose
WT: wild-type

## Acknowledgments

We would like to thank Dr. Toyoshi Fujimoto (Juntendo University) for kindly gifting the wild-type yeast (SEY6210) and Editage (www.editage.com) for English language editing.

## Author contributions

All authors contributed to the conception and design of this study. N. M., R. K., M. M., K. F., S. K., T. M., and A. F. performed material preparation, data collection, and analysis. A. F. wrote the first draft of the manuscript, and all authors commented on the previous versions of the manuscript. All the authors have read and approved the final version of the manuscript.

## Data availability

All data generated or analyzed during this study are included in this published article (and its Supplementary information files).

## Supplemental information

**Table S1.**
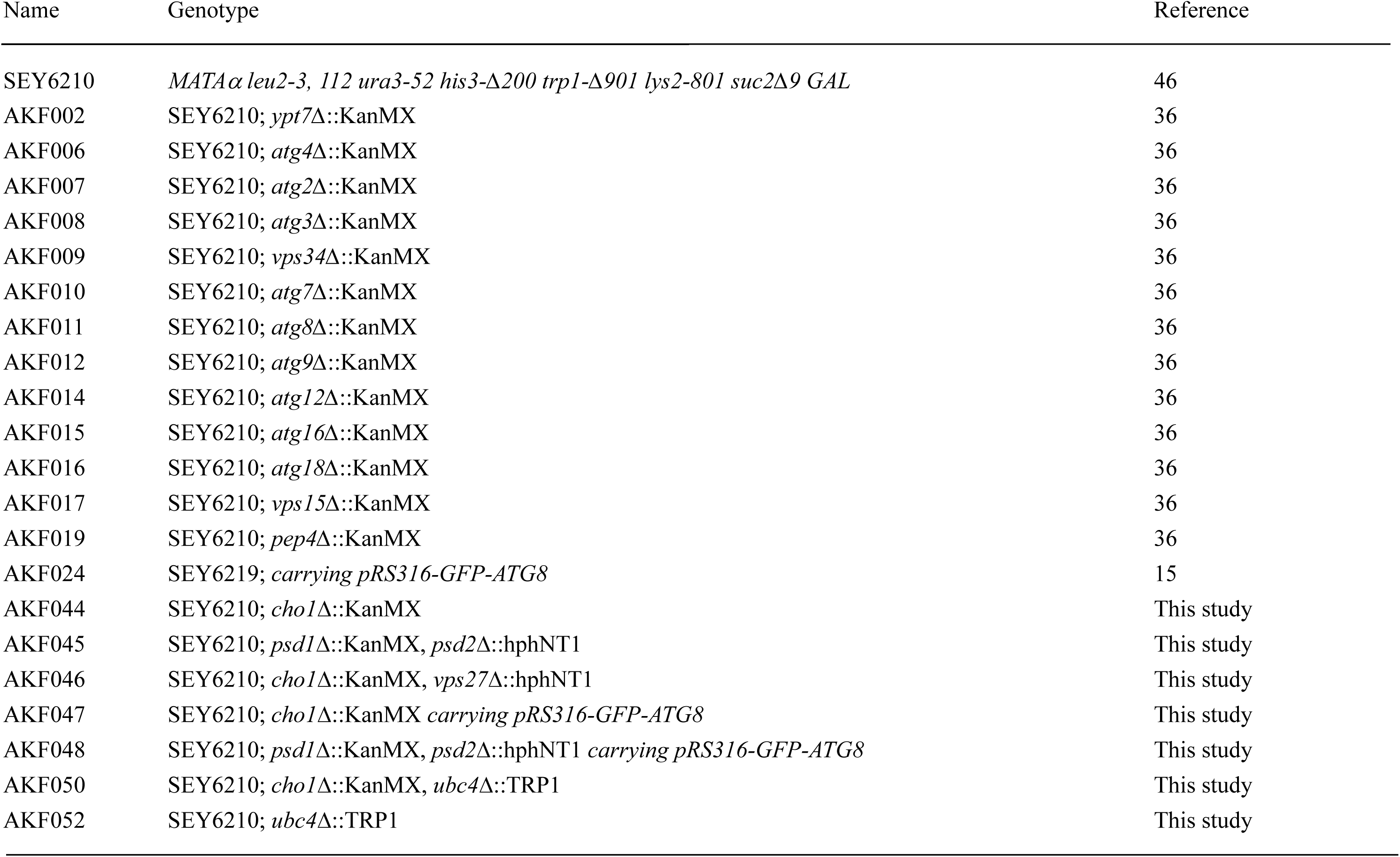
Yeast strains used in this study. Deletion mutants were generated using a standard PCR-based method.

**Figure S1.**
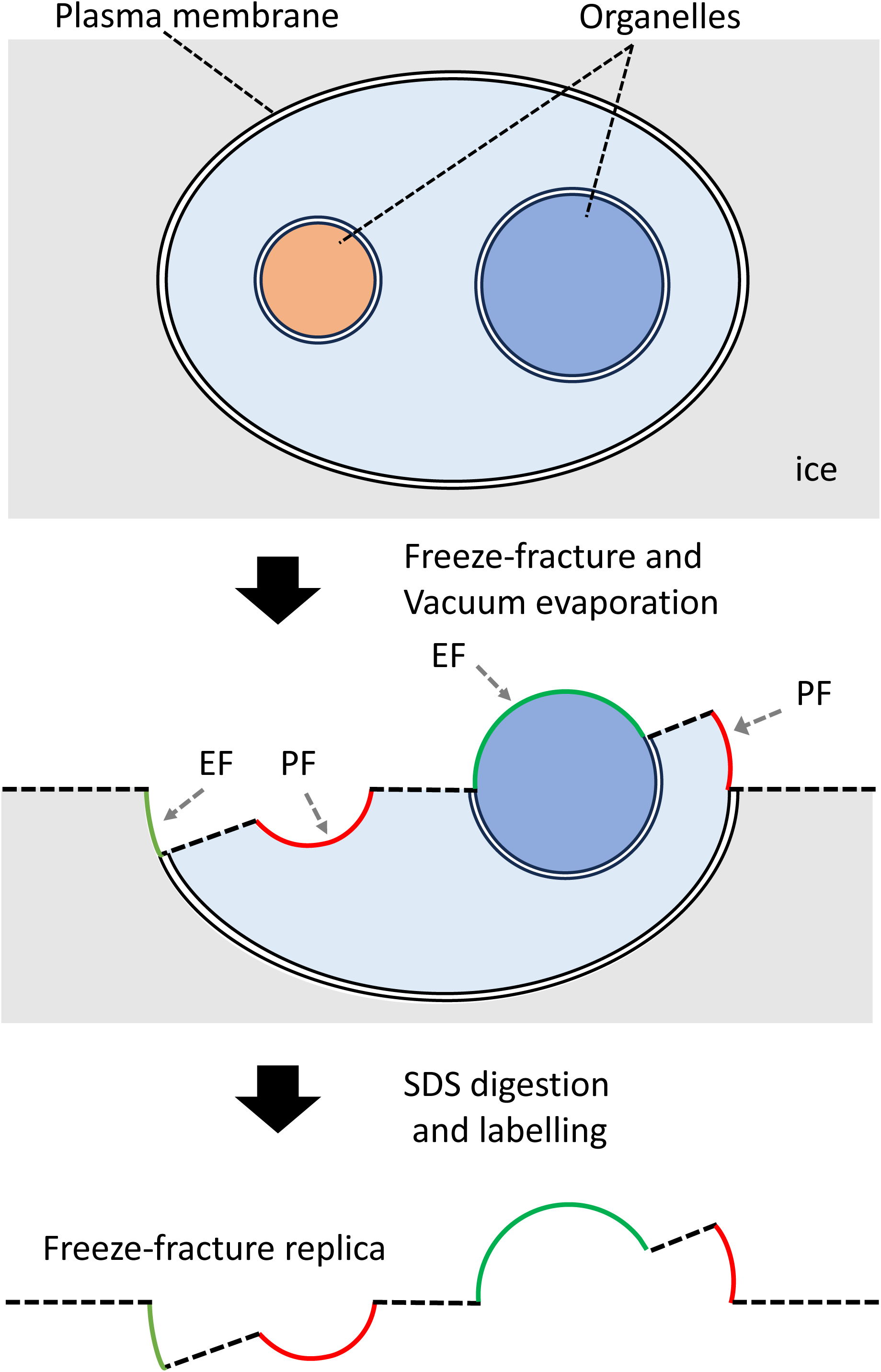
Outline of freeze-fracture EM. Replicas representing the outer (luminal) and inner (cytoplasmic) membrane leaflets are called the EF and the PF, respectively. Fracture planes passing through the cytoplasm and the extracellular space are shown by a dotted line.

**Figure S2.**
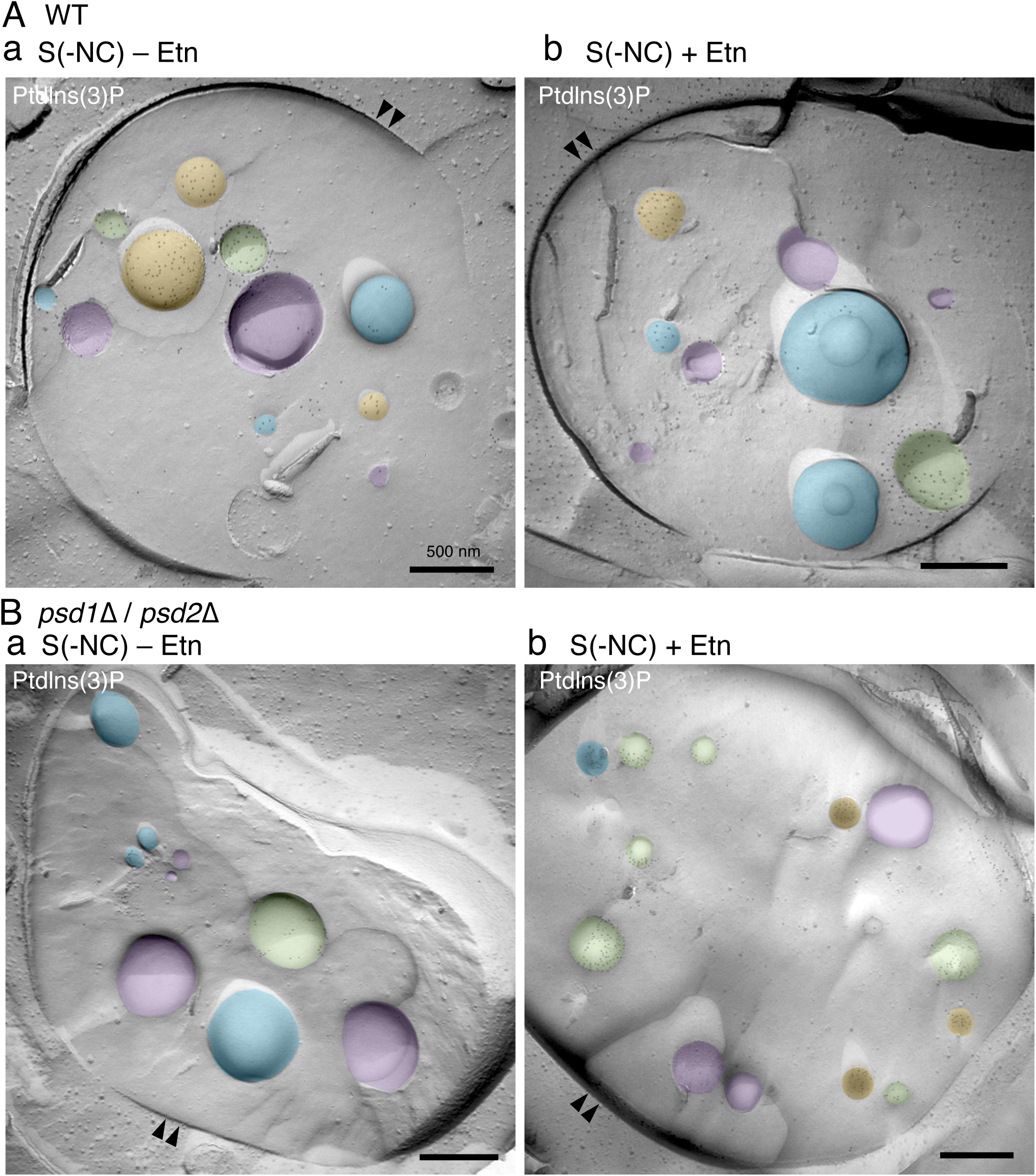
Macroautophagic body and microautophagic vesicle in the vacuole of WT and *psd1*Δ/*psd2*Δ yeast strains. WT yeast (A) and the *psd1*Δ/*psd2*Δ (B) mutant were starved in S(-NC) medium supplemented with PMSF for 5 h at 30 °C, both in the absence (a) and presence (b) of ethanolamine (Etn). In WT yeast (A) and the *psd1*Δ/*psd2*Δ (B) mutant, two types of vesicles were observed under both conditions: one with PtdIns(3)P labeling enriched in the external face (EF, green) of vesicles in the vacuoles and another with labeling in the peripheral face (PF, orange), but not the EF (purple). Double arrowheads, vacuole. Scale bar, 500 nm.

**Figure S3.**
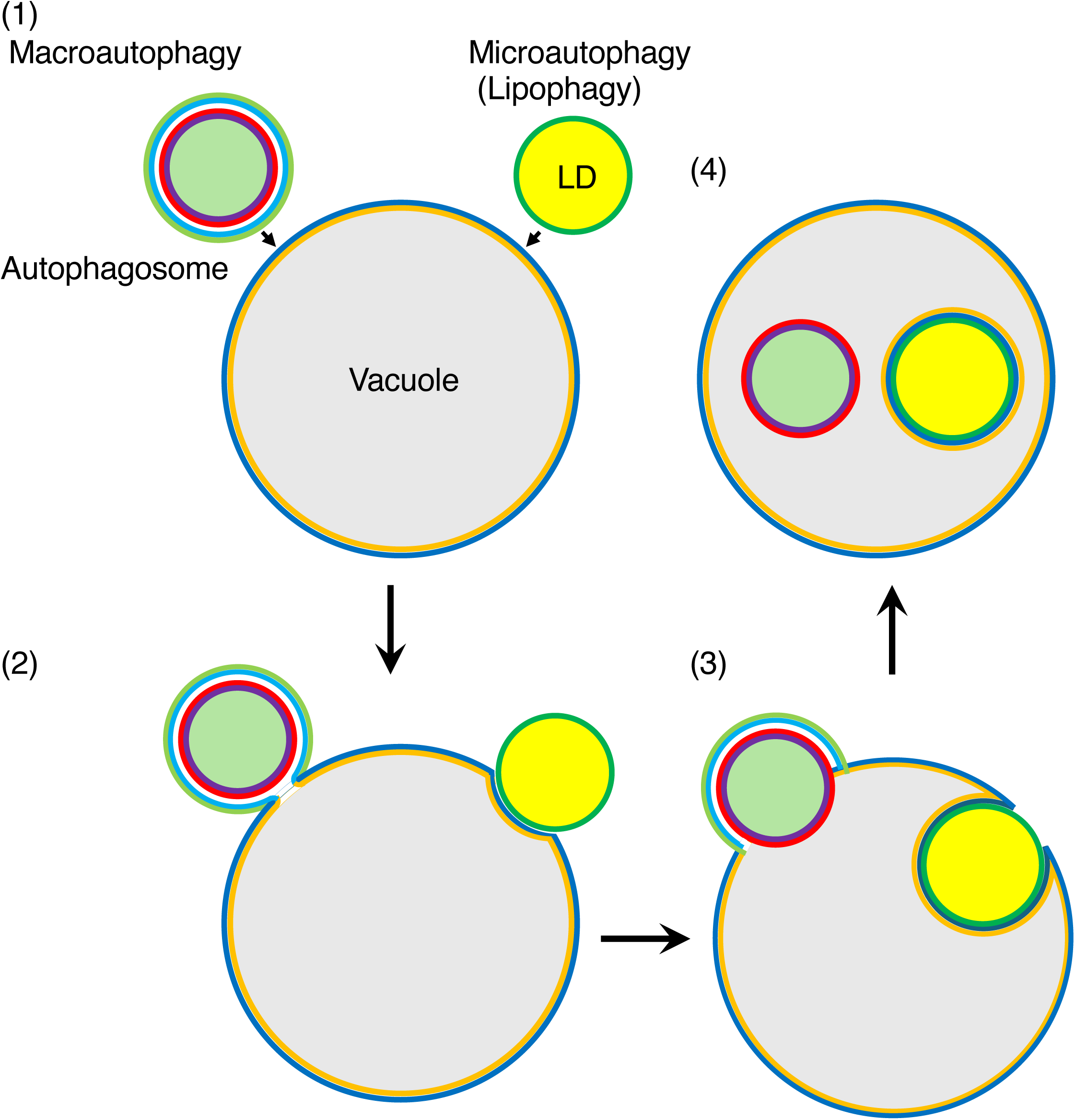
Forming processes of macroautophagic body and microautophagic vesicle in the vacuole. Diagram showing how the macroautophagic body and microautophagic vesicle form in the vacuole. LD, lipid droplet.

**Figure S4.**
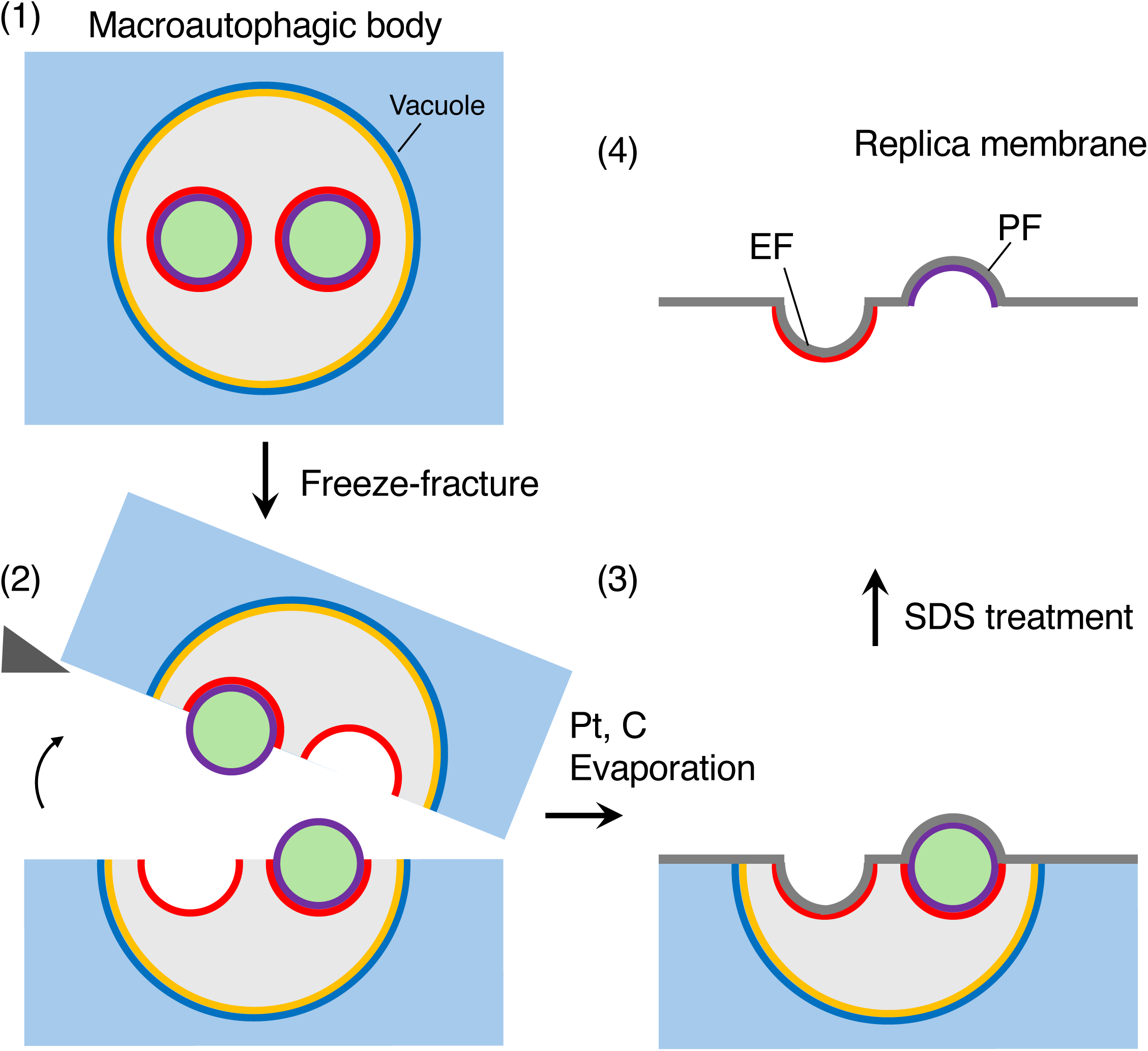
Outline of the quick-freezing and freeze-fracture labelling (QF-FRL) method of macroautophagic bodies in the vacuole. (1) Quick-freezing (QF): Live yeast cells are quickly frozen without ice crystal formation. High-pressure and metal contact freezing methods were used in the present study. In the high-pressure freezing method, samples are frozen using liquid nitrogen, and the nucleation and growth of ice crystals were reduced by the brief application of a pressure of 2100 bar. (2) Freeze-fracture: Frozen yeast cells are fractured at below − 130°C and under a high vacuum. Membranes are split into two leaflets, and the hydrophobic interface (i.e. the acyl chain side of the phospholipid monolayer) is exposed. (3) Vacuum evaporation: thin layers of carbon (C) and platinum (Pt) are deposited onto the hydrophobic interface of membranes and physically stabilize the molecules. Because platinum is evaporated at an oblique angle to the surface of the specimen (45° in the present study), protruding structures block the evaporating atoms to produce ‘shadows’ behind the structures. Areas deficient in the platinum deposition, therefore, appear to be electron-lucent under EM. Transmembrane proteins are observed as small bumps termed intra-membrane particles (IMPs). (4) SDS treatment. Specimens are thawed and treated with a SDS solution to dissolve materials other than the lipid monolayer and integral membrane proteins, which are in direct contact with the carbon and platinum layer. This makes membrane proteins and lipid head groups accessible for antibody labelling.

**Figure S5.**
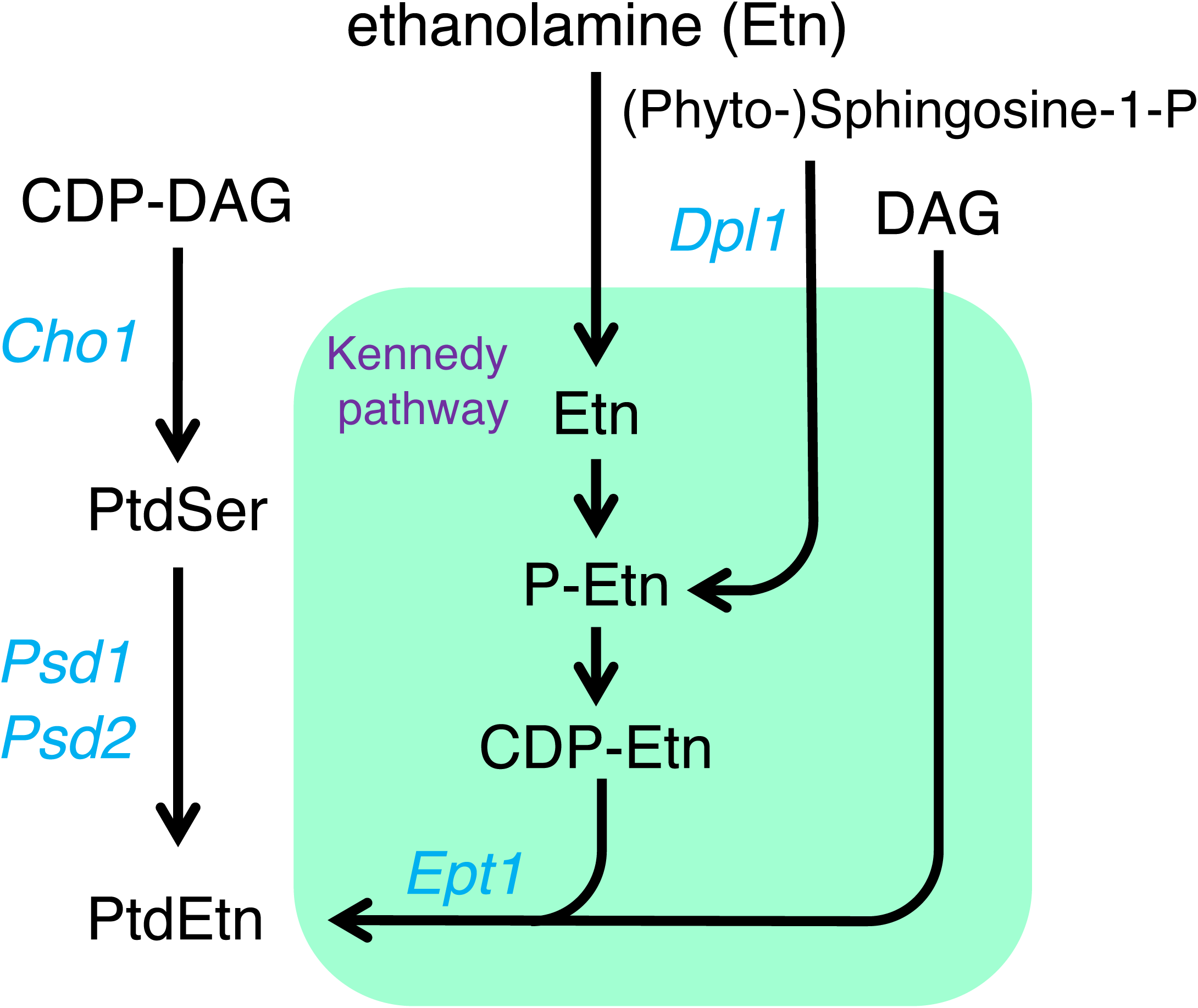
Pathways of PtdEtn biosynthesis. In *Saccharomyces cerevisiae*, PtdEtn is synthesized through three primary pathways: 1) the de novo or CDP-DAG pathway via mitochondrial Psd1, 2) the CDP-DAG pathway via Psd2, and 3) the CDP-ethanolamine branch of the Kennedy pathway.

**Fig. S6.**
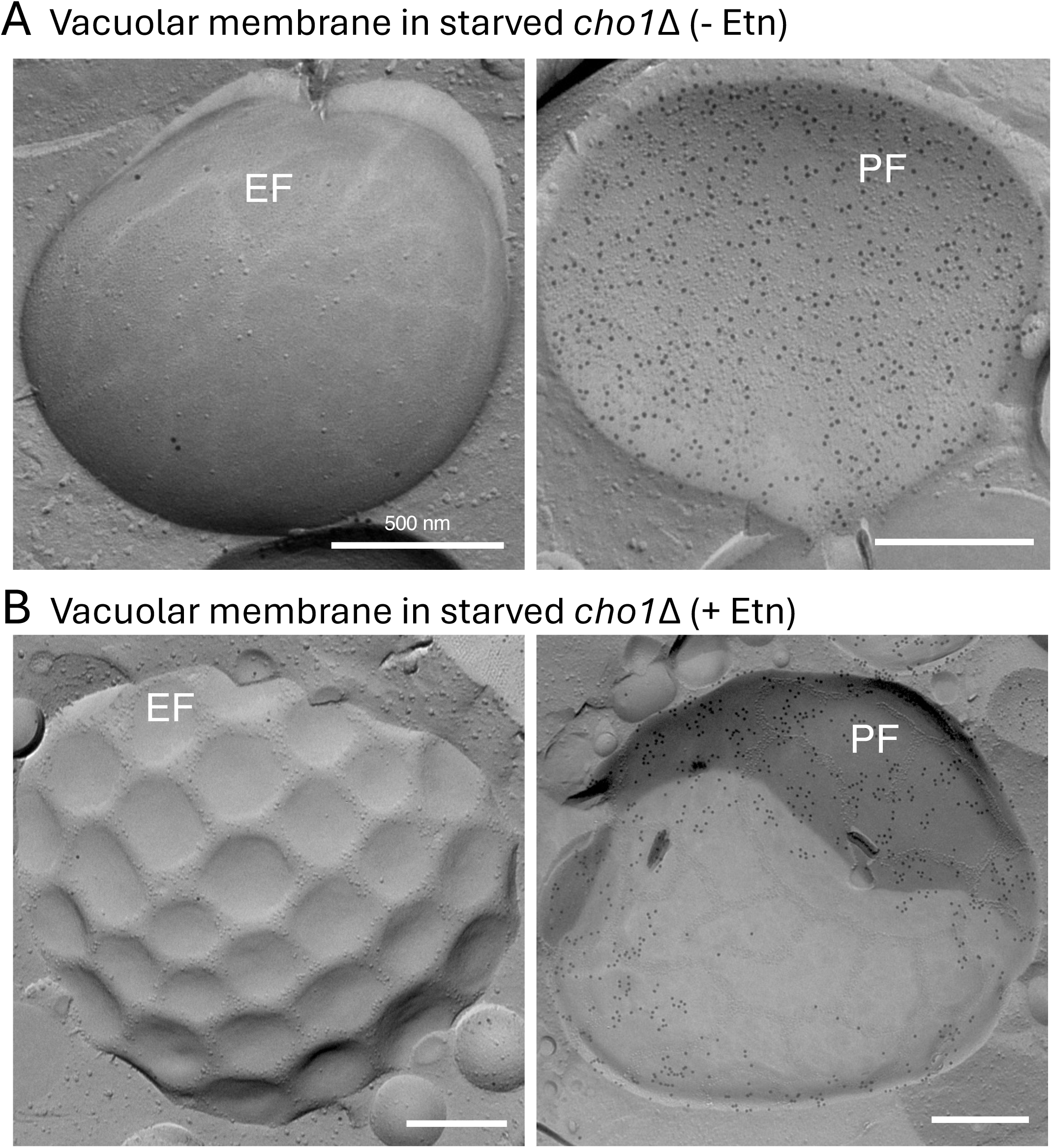
Geometric patterns of the vacuolar membrane in starved *cho1*Δ mutants with ethanolamine supplementation. The *cho1*Δ mutant was starved in S(-NC) medium for 5 h, either in the absence (A) or presence (B) of ethanolamine (Etn). In the absence of Etn, the vacuolar membrane displayed a typical round morphology in both the EF and PF fracture faces, as seen in (A). In contrast, in the presence of Etn, the vacuolar membrane exhibited distinct geometric patterns (B). PtdIns(3)P labeling was predominantly observed on the PF. Scale bar: 500 nm.

**Figure S7.**
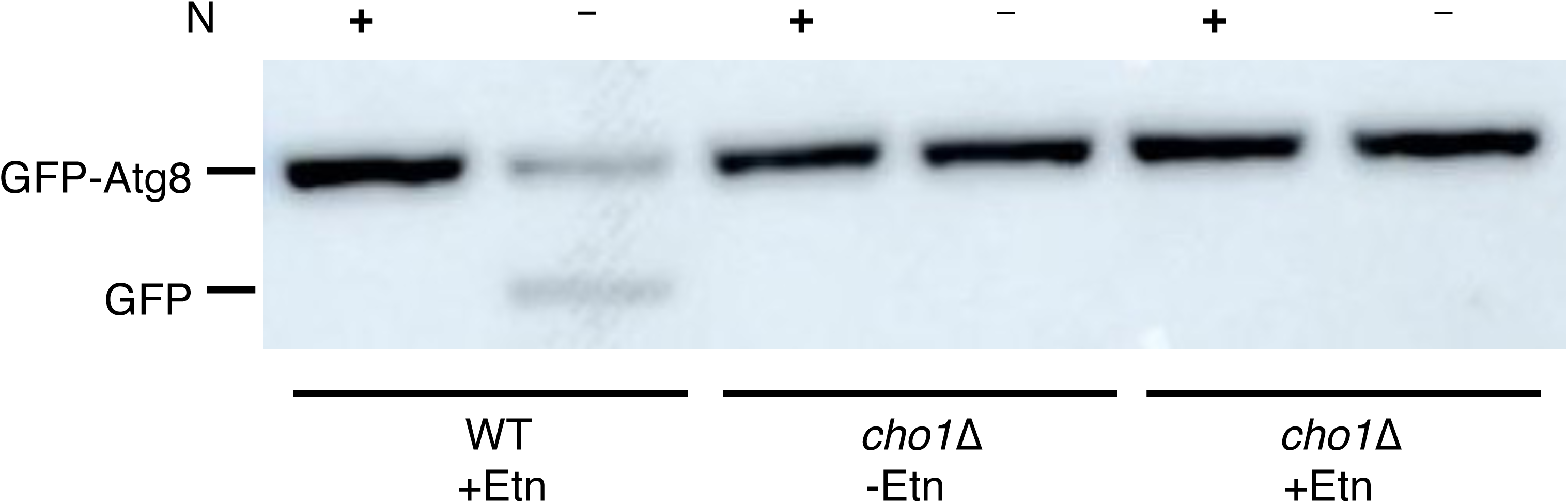
GFP-Atg8 processing assay under nitrogen-deficient SD(-N) conditions. Degradation of GFP-Atg8 via macroautophagy in the *cho1***Δ** mutant. Wild-type (WT) yeast and the *cho1***Δ** mutant, transformed with a plasmid expressing GFP-Atg8 under the control of the *CUP1* promoter, were cultured in SMD-Ura medium at 30°C to mid-log phase. The cultures were then divided into three aliquots: one was incubated in YPD medium for 4 hours at 30°C, while the other two were shifted to SD(-N) medium without PMSF and cultured for 4 h at 30°C, either in the presence or absence of 3 mM ethanolamine (Etn). Samples were collected before (+N) and after (-N) nitrogen starvation. Bands corresponding to full-length GFP-Atg8 and free GFP are shown. Impaired GFP-Atg8 degradation was observed in the *cho1***Δ** mutant regardless of Etn supplementation.

**Fig. S8.**
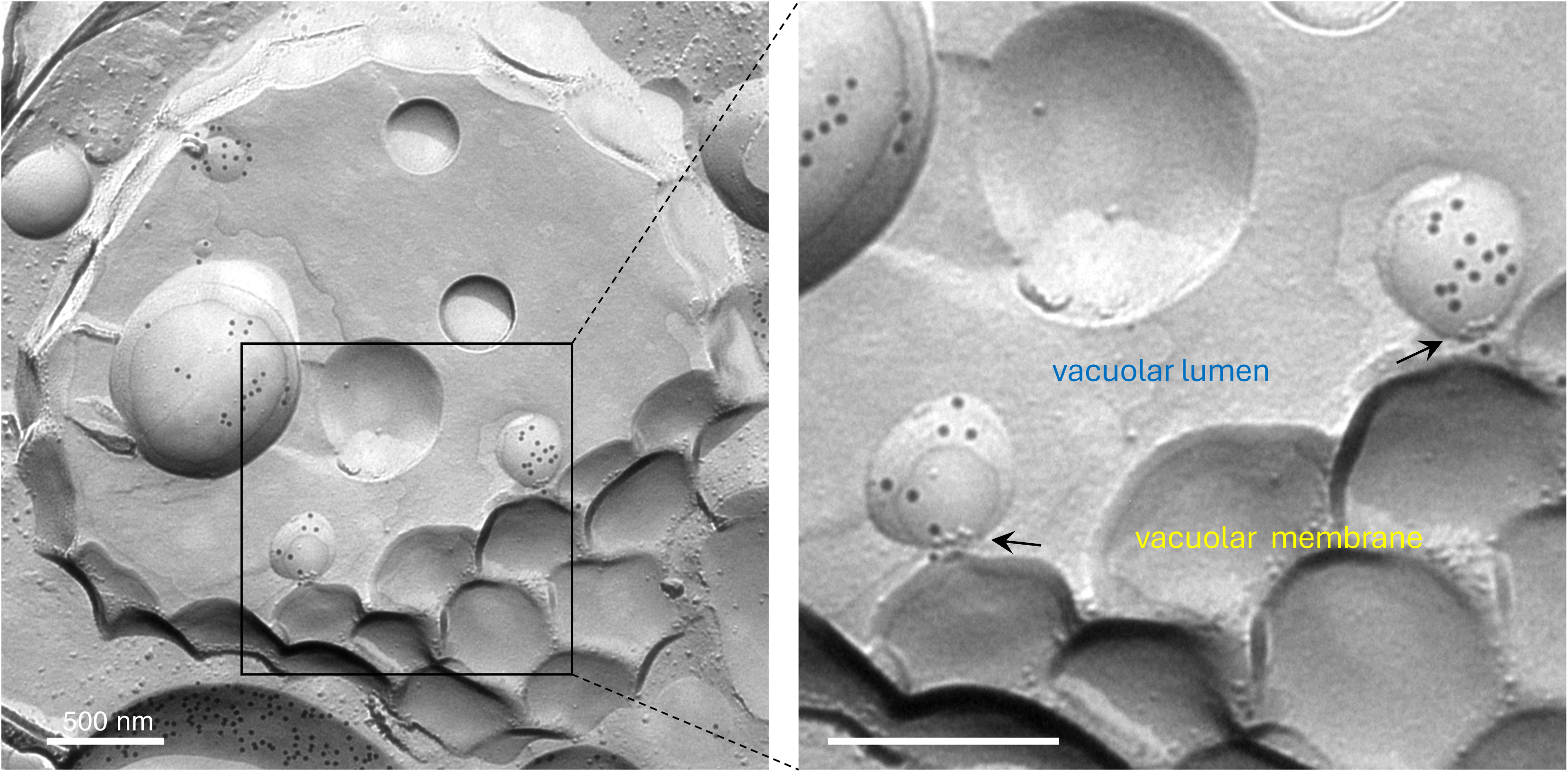
The vacuolar lumen contained vesicles associated with clusters of IMPs. The cho1Δ mutant was starved for 5 hr in S(-NC) supplemented with PMSF. A raft-like domain gave rise to a balloon-shaped projection extending into the vacuolar lumen. Clusters of IMPs appeared as a ring at the neck of this structure (indicated by arrows). The labelings of PtdIns(3)P were observed in the PF, but not EF, of the microautophagic vesicles.

**Figure S9.**
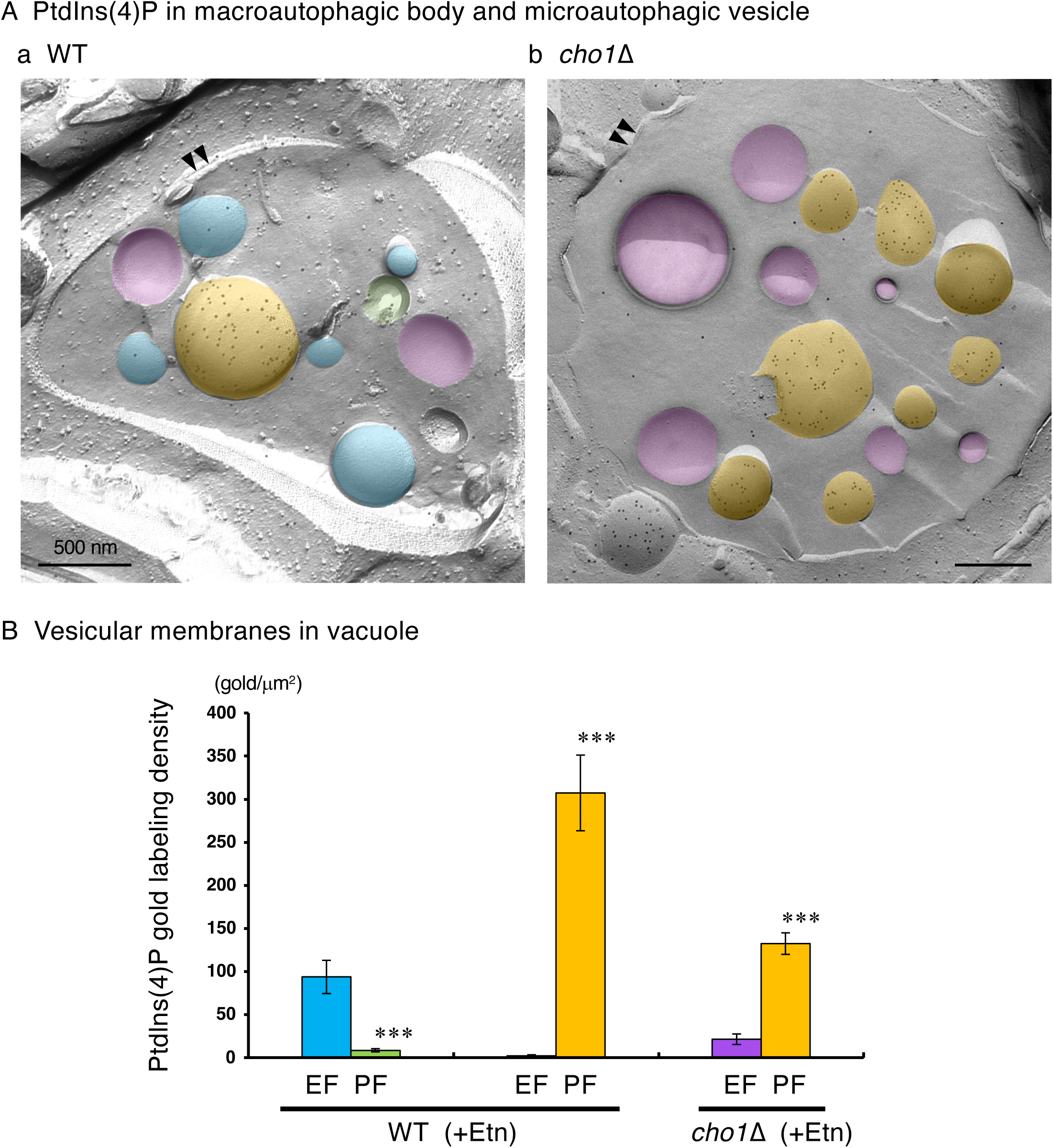
PtdIns(4)P labeling in vesicle membranes within the vacuole of starved WT and *cho1*Δ yeast. (A) Electron micrographs showing PtdIns(4)P labeling on the PF and EF of macroautophagic body membranes (blue and green) and microautophagic vesicle membranes (orange and purple) within the vacuole of WT and *cho1Δ* mutant yeast starved in S(-NC) supplemented with PMSF in the presence of ethanolamine (Etn). Scale bar: 500 nm. (B) Quantification of PtdIns(4)P labeling density on the PF of microautophagic vesicles and the EF of macroautophagic bodies in the vacuoles of starved WT and *cho1Δ* yeast. Labeling density on the PF of microautophagic vesicles and the EF of macroautophagic bodies was significantly higher than on the opposite membrane faces (EF and PF, respectively), indicating that the vesicles in the vacuole of starved *cho1Δ* cells are predominantly microautophagic in origin. Data represent mean ± SE from three independent experiments (>30 vesicles analyzed per group in each experiment). Statistical analysis was performed using a *t*-test: \*\*\**p* < 0.001.

**Figure S10.**
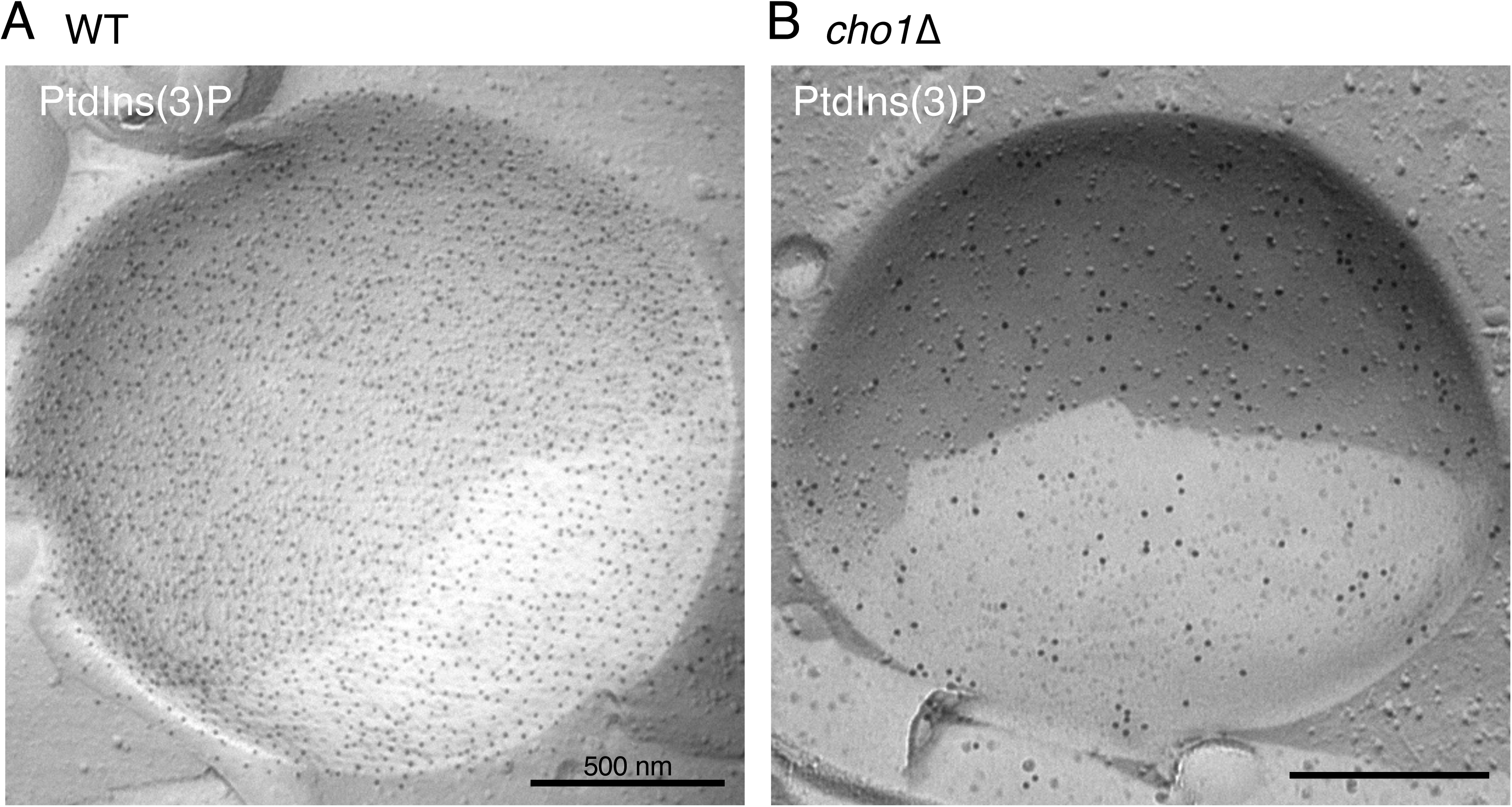
Decrease in the PtdIns(3)P labelings in the vacuolar membrane of *cho1*Δ mutant. Micrograph shows the PtdIns(3)P labelings on the PF in the vacuolar membranes of WT (A) and *cho1*Δ (B) strains starved in S(-NC) supplemented with PMSF in the presence of ethanolamine (Etn). The PtdIns(3)P labeling in the vacuolar membrane of *cho1*Δ mutant was much lower than that of WT yeast.

**Figure S11.**
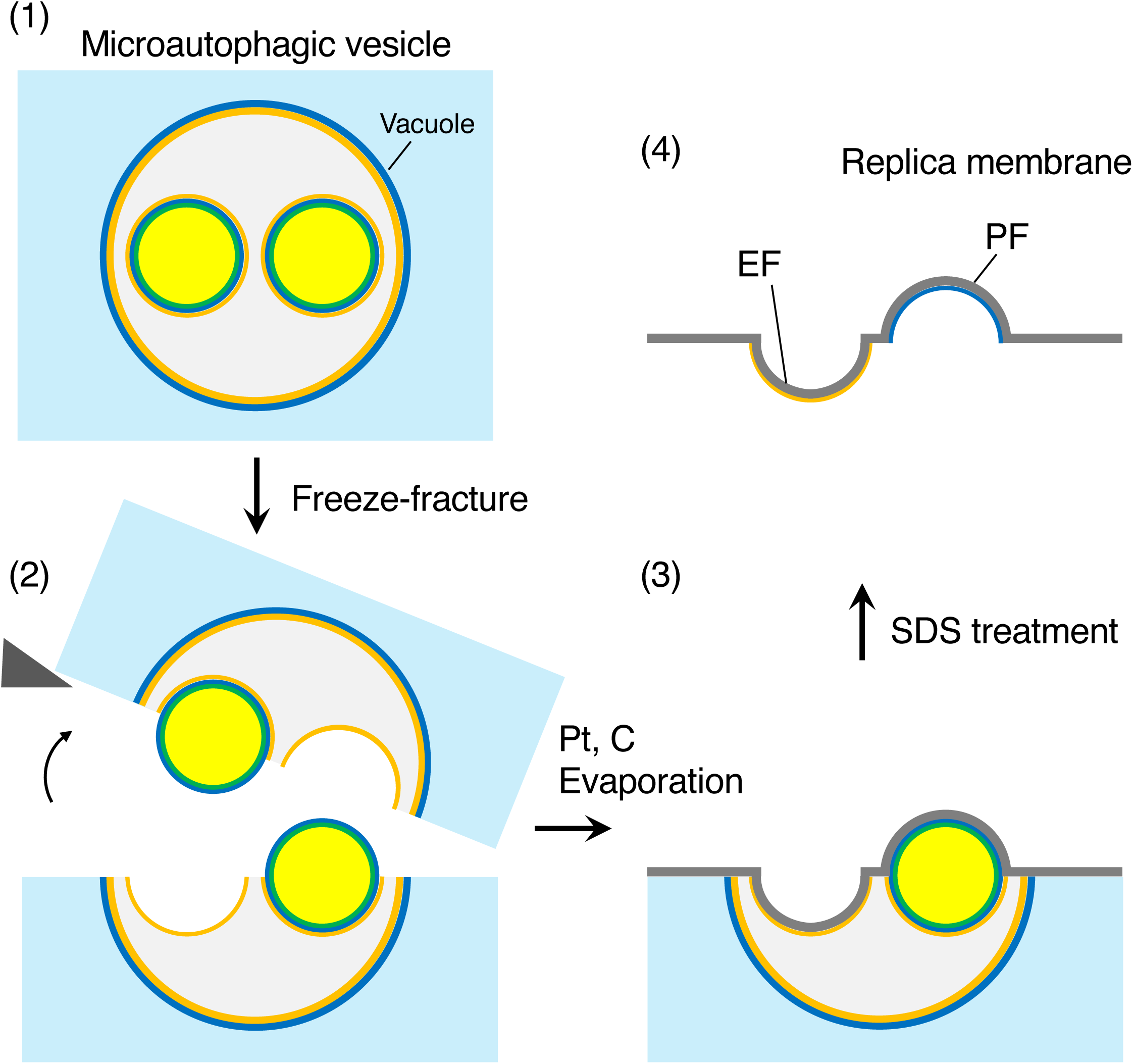
Outline of the QF-FRL method of microautophagic vesicles in the vacuole. The detail of the QF-FRL method is described in the figure legend of Fig. S4.

**Figure S12.**
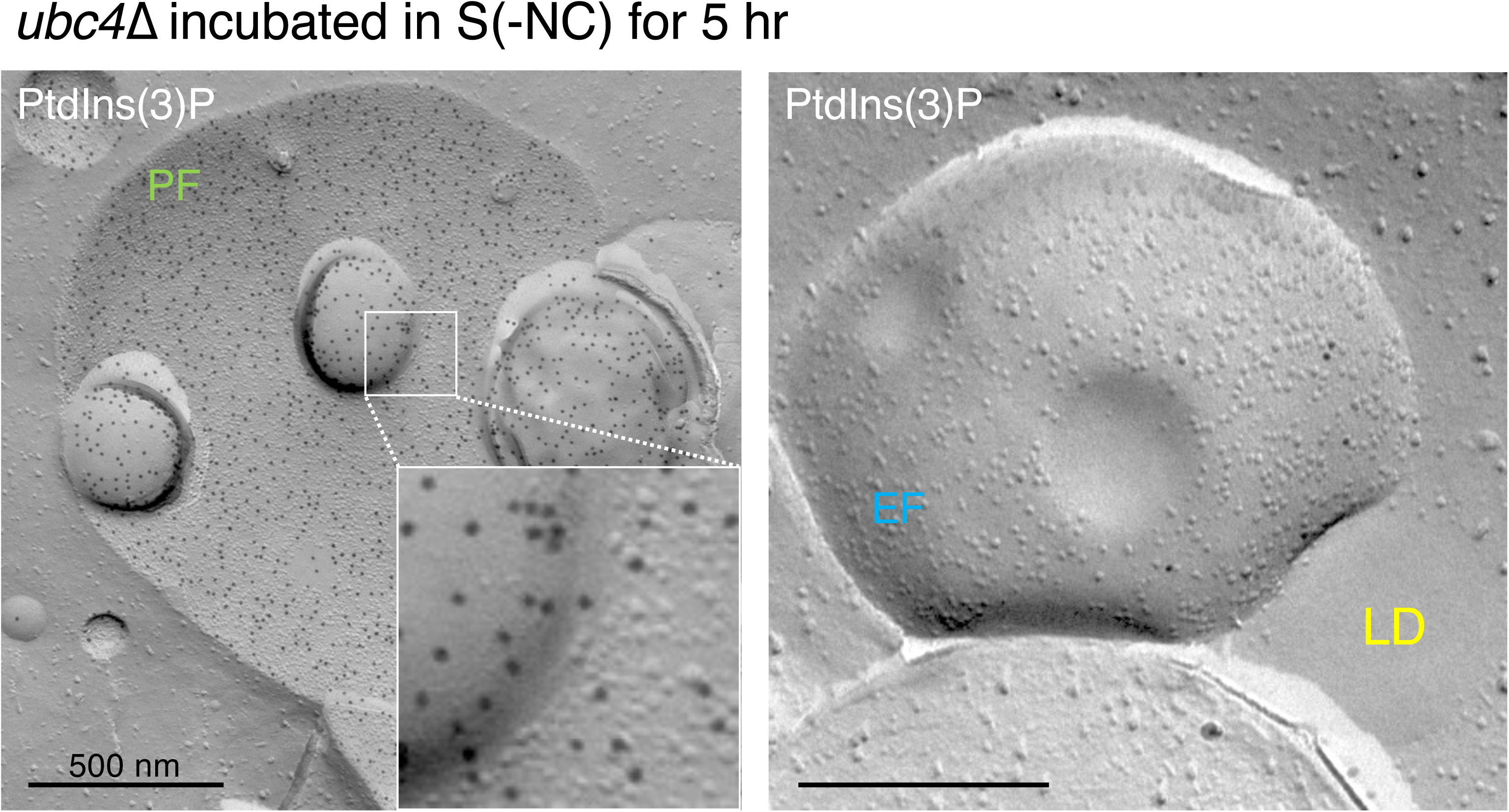
IMP-deficient raft-like domains in the vacuolar membrane of *ubc4*Δ mutant after starvation. The IMP-deficient raft-like domains were observed both in the PF (left) and the EF (right) of the vacuolar membrane in the starved *ubc4*Δ mutant supplemented with PMSF. The PF is densely labeled for PtdIns(3)P both in the IMP-rich and IMP-deficient domains. Scale bar: 500 nm.

**Figure S13.**
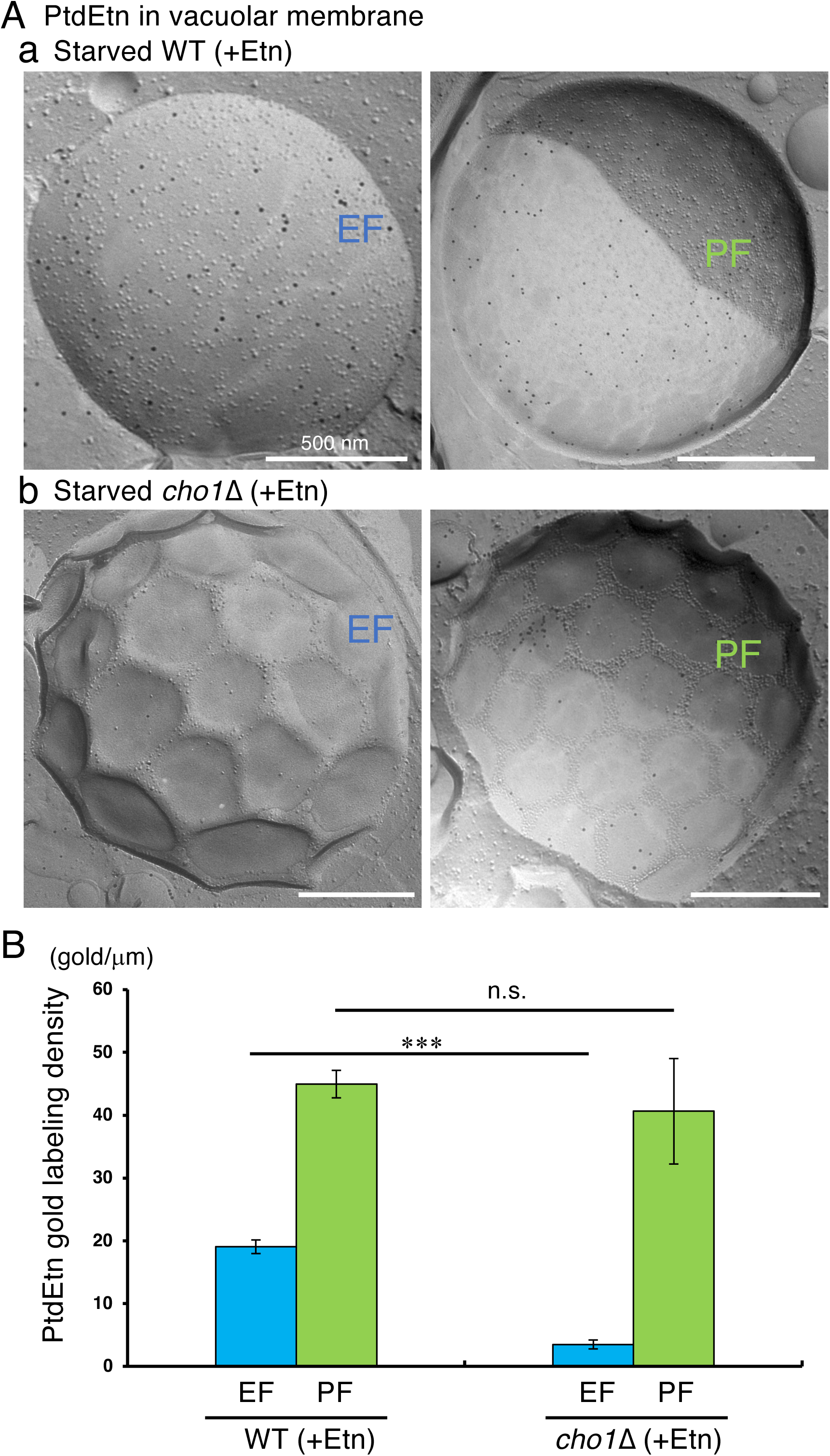
Distribution of PtdEtn in the vacuolar membrane of the starved WT and *cho1*Δ strain. (A) PtdEtn was localized in both the EF and PF leaflets in the vacuolar membrane of the starved WT yeast supplemented with PMSF in the presence of Etn in the medium, and the PtdEtn labeling density was much higher in the PF than that in the EF of WT (B). In contrast to the WT yeast, the PtdEtn labeling was detected on the PF, but not the EF, in the vacuolar membrane of the starved *cho1*Δ in the presence of Etn. Scale bar: 500 nm. Mean ± SE of three independent experiments (>30 vacuoles were counted in each experiment). The PtdEtn labeling of the *cho1*Δ mutant was compared to that of the WT yeast. *t-test*: n.s., not significant; *** *p* < 0.001.

**Fig. S14.**
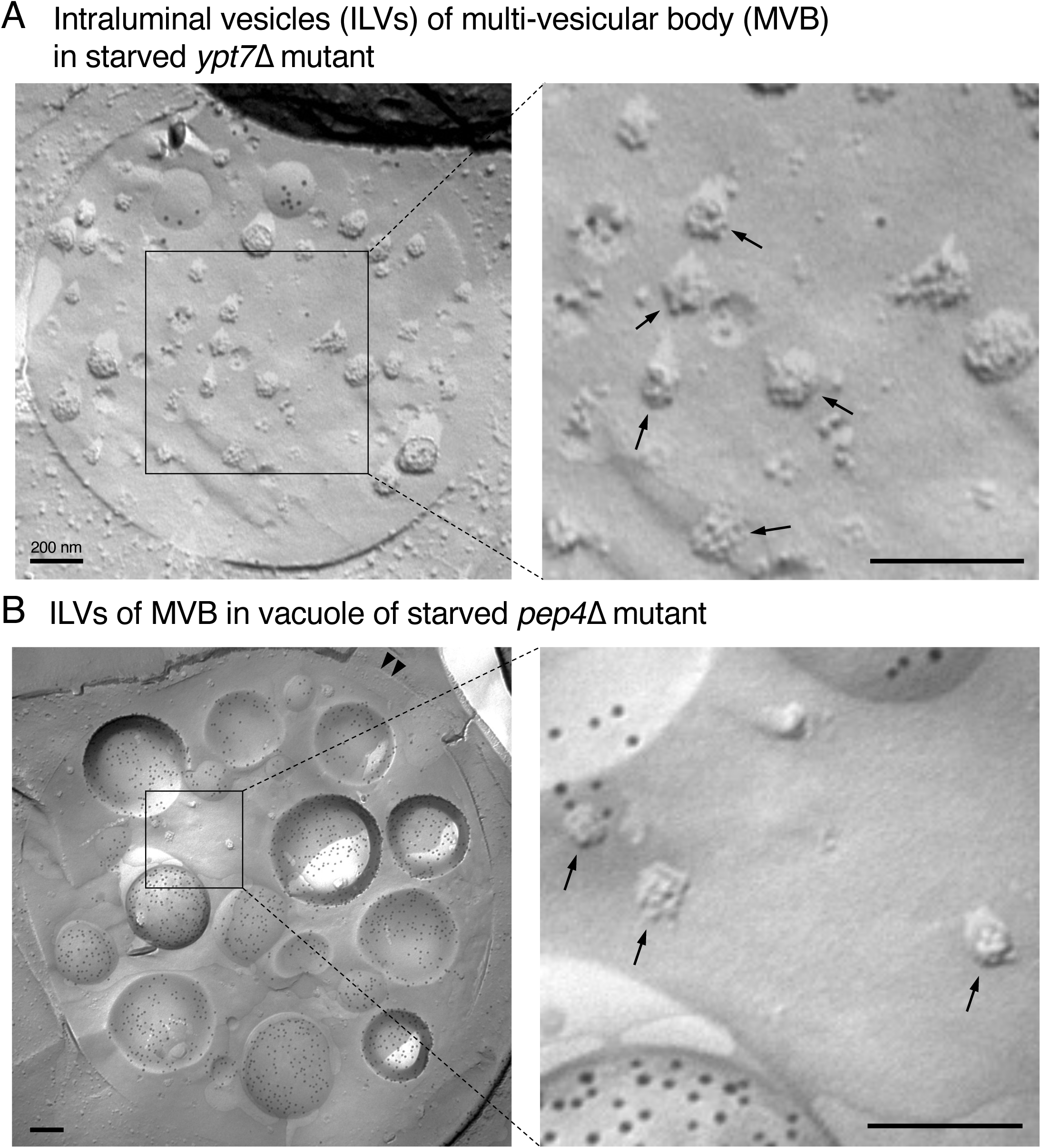
Intraluminal vesicles (ILVs) within the multivesicular bodies (MVBs) of starved *ypt7*Δ mutants and in the vacuolar lumen of starved *pep4*Δ mutants. The *ypt7*Δ and *pep4*Δ mutants were subjected to starvation in S(-NC) medium without PMSF at 30°C for 5 hours. (A) In starved *ypt7*Δ mutants, intraluminal vesicles (ILVs), appearing as small, irregularly contoured structures (indicated by arrows in the right panel), were observed within the multivesicular bodies (MVBs) of endosomes using the freeze-fracture EM method. (B) In starved *pep4*Δ cells, ILVs along with accumulated macroautophagic bodies labeled for PtdSer on both the EF and PF were detected within the vacuole. The morphology of these rugged structures was indistinguishable from that of ILVs observed in the endosomal MVBs of *ypt7*Δ mutants (see also Figure 6—figure supplement 1 in [10]).

## References

[1] N. Mizushima, M. Komatsu, Autophagy: renovation of cells and tissues, Cell, 147 (2011) 728–741.

[2] Z. Xie, D.J. Klionsky, Autophagosome formation: core machinery and adaptations, Nat Cell Biol, 9 (2007) 1102–1109.

[3] D. Mijaljica, M. Prescott, R.J. Devenish, Microautophagy in mammalian cells: revisiting a 40-year-old conundrum, Autophagy, 7 (2011) 673–682.

[4] C. De Duve, R. Wattiaux, Functions of lysosomes, Annu Rev Physiol, 28 (1966) 435–492.

[5] D.L. Tuttle, W.A. Dunn, Jr., Divergent modes of autophagy in the methylotrophic yeast Pichia pastoris, J Cell Sci, 108 (Pt 1) (1995) 25–35.

[6] T. Osawa, T. Kotani, T. Kawaoka, E. Hirata, K. Suzuki, H. Nakatogawa, Y. Ohsumi, N.N. Noda, Atg2 mediates direct lipid transfer between membranes for autophagosome formation, Nat Struct Mol Biol, 26 (2019) 281–288.

[7] S. Maeda, H. Yamamoto, L.N. Kinch, C.M. Garza, S. Takahashi, C. Otomo, N.V. Grishin, S. Forli, N. Mizushima, T. Otomo, Structure, lipid scrambling activity and role in autophagosome formation of ATG9A, Nat Struct Mol Biol, 27 (2020) 1194–1201.

[8] K. Matoba, T. Kotani, A. Tsutsumi, T. Tsuji, T. Mori, D. Noshiro, Y. Sugita, N. Nomura, S. Iwata, Y. Ohsumi, T. Fujimoto, H. Nakatogawa, M. Kikkawa, N.N. Noda, Atg9 is a lipid scramblase that mediates autophagosomal membrane expansion, Nat Struct Mol Biol, 27 (2020) 1185–1193.

[9] Y. Kurokawa, R. Konishi, A. Yoshida, K. Tomioku, K. Tanabe, A. Fujita, Microautophagy in the yeast vacuole depends on the activities of phosphatidylinositol 4-kinases, Stt4p and Pik1p, Biochim Biophys Acta Biomembr, 1862 (2020) 183416.

[10] T. Tsuji, M. Fujimoto, T. Tatematsu, J. Cheng, M. Orii, S. Takatori, T. Fujimoto, Niemann-Pick type C proteins promote microautophagy by expanding raft-like membrane domains in the yeast vacuole, Elife, 6 (2017).

[11] S. Maeda, C. Otomo, T. Otomo, The autophagic membrane tether ATG2A transfers lipids between membranes, Elife, 8 (2019).

[12] K. Tsuboyama, I. Koyama-Honda, Y. Sakamaki, M. Koike, H. Morishita, N. Mizushima, The ATG conjugation systems are important for degradation of the inner autophagosomal membrane, Science, 354 (2016) 1036–1041.

[13] Y. Kurokawa, A. Yoshida, E. Fujii, K. Tomioku, H. Hayashi, K. Tanabe, A. Fujita, Phosphatidylinositol 4-phosphate on Rab7-positive autophagosomes revealed by the freeze-fracture replica labeling, Traffic, 20 (2019) 82–95.

[14] Y. Yamakuchi, Y. Kurokawa, R. Konishi, K. Fukuda, T. Masatani, A. Fujita, Selective increment of phosphatidylserine on the autophagic body membrane in the yeast vacuole, FEBS Lett, 595 (2021) 2197–2207.

[15] M. Muramoto, N. Mineoka, K. Fukuda, S. Kuriyama, T. Masatani, A. Fujita, Coordinated regulation of phosphatidylinositol 4-phosphate and phosphatidylserine levels by Osh4p and Osh5p is an essential regulatory mechanism in autophagy, Biochim Biophys Acta Biomembr, 1866 (2024) 184308.

[16] J.H. Hurley, T. Meyer, Subcellular targeting by membrane lipids, Curr Opin Cell Biol, 13 (2001) 146–152.

[17] P. Varnai, T. Balla, Visualization of phosphoinositides that bind pleckstrin homology domains: calcium– and agonist-induced dynamic changes and relationship to myo-[3H]inositol-labeled phosphoinositide pools, J Cell Biol, 143 (1998) 501–510.

[18] H. Bisio, A. Krishnan, J.B. Marq, D. Soldati-Favre, Toxoplasma gondii phosphatidylserine flippase complex ATP2B-CDC50.4 critically participates in microneme exocytosis, PLoS Pathog, 18 (2022) e1010438.

[19] T.E. Rusten, H. Stenmark, Analyzing phosphoinositides and their interacting proteins, Nat Methods, 3 (2006) 251–258.

[20] G.R. Hammond, T. Balla, Polyphosphoinositide binding domains: Key to inositol lipid biology, Biochim Biophys Acta, 1851 (2015) 746–758.

[21] O.V. Vieira, R.E. Harrison, C.C. Scott, H. Stenmark, D. Alexander, J. Liu, J. Gruenberg, A.D. Schreiber, S. Grinstein, Acquisition of Hrs, an essential component of phagosomal maturation, is impaired by mycobacteria, Mol Cell Biol, 24 (2004) 4593–4604.

[22] S.A. Watt, G. Kular, I.N. Fleming, C.P. Downes, J.M. Lucocq, Subcellular localization of phosphatidylinositol 4,5-bisphosphate using the pleckstrin homology domain of phospholipase C delta1, Biochem J, 363 (2002) 657–666.

[23] J. van Rheenen, E.M. Achame, H. Janssen, J. Calafat, K. Jalink, PIP2 signaling in lipid domains: a critical re-evaluation, EMBO J, 24 (2005) 1664–1673.

[24] G.D. Fairn, N.L. Schieber, N. Ariotti, S. Murphy, L. Kuerschner, R.I. Webb, S. Grinstein, R.G. Parton, High-resolution mapping reveals topologically distinct cellular pools of phosphatidylserine, J Cell Biol, 194 (2011) 257–275.

[25] A. Fujita, J. Cheng, T. Fujimoto, Quantitative electron microscopy for the nanoscale analysis of membrane lipid distribution, Nat Protoc, 5 (2010) 661–669.

[26] A. Fujita, J. Cheng, M. Hirakawa, K. Furukawa, S. Kusunoki, T. Fujimoto, Gangliosides GM1 and GM3 in the living cell membrane form clusters susceptible to cholesterol depletion and chilling, Mol Biol Cell, 18 (2007) 2112–2122.

[27] A. Fujita, J. Cheng, K. Tauchi-Sato, T. Takenawa, T. Fujimoto, A distinct pool of phosphatidylinositol 4,5-bisphosphate in caveolae revealed by a nanoscale labeling technique, Proc Natl Acad Sci U S A, 106 (2009) 9256–9261.

[28] A. Yoshida, M. Shigekuni, K. Tanabe, A. Fujita, Nanoscale analysis reveals agonist-sensitive and heterogeneous pools of phosphatidylinositol 4-phosphate in the plasma membrane, Biochim Biophys Acta, 1858 (2016) 1298–1305.

[29] J.S. Robinson, D.J. Klionsky, L.M. Banta, S.D. Emr, Protein sorting in Saccharomyces cerevisiae: isolation of mutants defective in the delivery and processing of multiple vacuolar hydrolases, Mol Cell Biol, 8 (1988) 4936–4948.

[30] U. Guldener, S. Heck, T. Fielder, J. Beinhauer, J.H. Hegemann, A new efficient gene disruption cassette for repeated use in budding yeast, Nucleic Acids Res, 24 (1996) 2519–2524.

[31] K. Suzuki, T. Kirisako, Y. Kamada, N. Mizushima, T. Noda, Y. Ohsumi, The pre-autophagosomal structure organized by concerted functions of APG genes is essential for autophagosome formation, EMBO J, 20 (2001) 5971–5981.

[32] H. Cheong, D.J. Klionsky, Biochemical methods to monitor autophagy-related processes in yeast, Methods Enzymol, 451 (2008) 1–26.

[33] T. Fujimoto, K. Fujimoto, Metal sandwich method to quick-freeze monolayer cultured cells for freeze-fracture, J Histochem Cytochem, 45 (1997) 595–598.

[34] J. Cheng, A. Fujita, H. Yamamoto, T. Tatematsu, S. Kakuta, K. Obara, Y. Ohsumi, T. Fujimoto, Yeast and mammalian autophagosomes exhibit distinct phosphatidylinositol 3-phosphate asymmetries, Nat Commun, 5 (2014) 3207.

[35] K. Obara, T. Noda, K. Niimi, Y. Ohsumi, Transport of phosphatidylinositol 3-phosphate into the vacuole via autophagic membranes in Saccharomyces cerevisiae, Genes Cells, 13 (2008) 537–547.

[36] Y. Kurokawa, R. Konishi, A. Yoshida, K. Tomioku, T. Futagami, H. Tamaki, K. Tanabe, A. Fujita, Essential and distinct roles of phosphatidylinositol 4-kinases, Pik1p and Stt4p, in yeast autophagy, Biochim Biophys Acta Mol Cell Biol Lipids, 1864 (2019) 1214–1225.

[37] M. Baba, M. Osumi, Y. Ohsumi, Analysis of the membrane structures involved in autophagy in yeast by freeze-replica method, Cell Struct Funct, 20 (1995) 465–471.

[38] A. Adachi, M. Koizumi, Y. Ohsumi, Autophagy induction under carbon starvation conditions is negatively regulated by carbon catabolite repression, J Biol Chem, 292 (2017) 19905–19918.

[39] T. Nishimura, N. Tamura, N. Kono, Y. Shimanaka, H. Arai, H. Yamamoto, N. Mizushima, Autophagosome formation is initiated at phosphatidylinositol synthase-enriched ER subdomains, EMBO J, 36 (2017) 1719–1735.

[40] A. Kihara, T. Noda, N. Ishihara, Y. Ohsumi, Two distinct Vps34 phosphatidylinositol 3-kinase complexes function in autophagy and carboxypeptidase Y sorting in Saccharomyces cerevisiae, J Cell Biol, 152 (2001) 519–530.

[41] K. Wang, Z. Yang, X. Liu, K. Mao, U. Nair, D.J. Klionsky, Phosphatidylinositol 4-kinases are required for autophagic membrane trafficking, J Biol Chem, 287 (2012) 37964–37972.

[42] Y. Ichimura, T. Kirisako, T. Takao, Y. Satomi, Y. Shimonishi, N. Ishihara, N. Mizushima, I. Tanida, E. Kominami, M. Ohsumi, T. Noda, Y. Ohsumi, A ubiquitin-like system mediates protein lipidation, Nature, 408 (2000) 488–492.

[43] Y. Kabeya, N. Mizushima, T. Ueno, A. Yamamoto, T. Kirisako, T. Noda, E. Kominami, Y. Ohsumi, T. Yoshimori, LC3, a mammalian homologue of yeast Apg8p, is localized in autophagosome membranes after processing, EMBO J, 19 (2000) 5720–5728.

[44] C.J. Clancey, S.C. Chang, W. Dowhan, Cloning of a gene (PSD1) encoding phosphatidylserine decarboxylase from Saccharomyces cerevisiae by complementation of an Escherichia coli mutant, J Biol Chem, 268 (1993) 24580–24590.

[45] P.J. Trotter, J. Pedretti, D.R. Voelker, Phosphatidylserine decarboxylase from Saccharomyces cerevisiae. Isolation of mutants, cloning of the gene, and creation of a null allele, J Biol Chem, 268 (1993) 21416–21424.

[46] P.J. Trotter, J. Pedretti, R. Yates, D.R. Voelker, Phosphatidylserine decarboxylase 2 of Saccharomyces cerevisiae. Cloning and mapping of the gene, heterologous expression, and creation of the null allele, J Biol Chem, 270 (1995) 6071–6080.

[47] S.A. Henry, S.D. Kohlwein, G.M. Carman, Metabolism and regulation of glycerolipids in the yeast Saccharomyces cerevisiae, Genetics, 190 (2012) 317–349.

[48] E. Calzada, O. Onguka, S.M. Claypool, Phosphatidylethanolamine Metabolism in Health and Disease, Int Rev Cell Mol Biol, 321 (2016) 29–88.

[49] P. Vigie, E. Cougouilles, I. Bhatia-Kissova, B. Salin, C. Blancard, N. Camougrand, The mitochondrial phosphatidylserine decarboxylase Psd1 is involved in nitrogen starvation-induced mitophagy in yeast, J Cell Sci, 132 (2019).

[50] C.H. Moeller, W.W. Thomson, An ultrastructural study of the yeast tomoplast during the shift from exponential to stationary phase, J Ultrastruct Res, 68 (1979) 28–37.

[51] A. Toulmay, W.A. Prinz, Direct imaging reveals stable, micrometer-scale lipid domains that segregate proteins in live cells, J Cell Biol, 202 (2013) 35–44.

[52] C.W. Wang, Y.H. Miao, Y.S. Chang, A sterol-enriched vacuolar microdomain mediates stationary phase lipophagy in budding yeast, J Cell Biol, 206 (2014) 357–366.

[53] R.C. Piper, A.A. Cooper, H. Yang, T.H. Stevens, VPS27 controls vacuolar and endocytic traffic through a prevacuolar compartment in Saccharomyces cerevisiae, J Cell Biol, 131 (1995) 603–617.

[54] M. Oku, Y. Maeda, Y. Kagohashi, T. Kondo, M. Yamada, T. Fujimoto, Y. Sakai, Evidence for ESCRT– and clathrin-dependent microautophagy, J Cell Biol, 216 (2017) 3263–3274.

[55] S. Morshed, M.N. Tasnin, T. Ushimaru, ESCRT machinery plays a role in microautophagy in yeast, BMC Mol Cell Biol, 21 (2020) 70.

[56] J.I. Sakamaki, K.L. Ode, Y. Kurikawa, H.R. Ueda, H. Yamamoto, N. Mizushima, Ubiquitination of phosphatidylethanolamine in organellar membranes, Mol Cell, 82 (2022) 3677–3692 e3611.

[57] T. van Zutphen, V. Todde, R. de Boer, M. Kreim, H.F. Hofbauer, H. Wolinski, M. Veenhuis, I.J. van der Klei, S.D. Kohlwein, Lipid droplet autophagy in the yeast Saccharomyces cerevisiae, Mol Biol Cell, 25 (2014) 290–301.

[58] G.M. Carman, G.S. Han, Regulation of phospholipid synthesis in the yeast Saccharomyces cerevisiae, Annu Rev Biochem, 80 (2011) 859–883.

[59] K.D. Atkinson, B. Jensen, A.I. Kolat, E.M. Storm, S.A. Henry, S. Fogel, Yeast mutants auxotrophic for choline or ethanolamine, J Bacteriol, 141 (1980) 558–564.

[60] J.I. Nikawa, S. Yamashita, Characterization of phosphatidylserine synthase from Saccharomyces cerevisiae and a mutant defective in the enzyme, Biochim Biophys Acta, 665 (1981) 420–426.

[61] Y. Kabeya, N. Mizushima, A. Yamamoto, S. Oshitani-Okamoto, Y. Ohsumi, T. Yoshimori, LC3, GABARAP and GATE16 localize to autophagosomal membrane depending on form-II formation, J Cell Sci, 117 (2004) 2805–2812.

[62] W.M. Henne, N.J. Buchkovich, S.D. Emr, The ESCRT pathway, Dev Cell, 21 (2011) 77–91.

[63] W. Seufert, S. Jentsch, Ubiquitin-conjugating enzymes UBC4 and UBC5 mediate selective degradation of short-lived and abnormal proteins, EMBO J, 9 (1990) 543–550.

[64] S.C. Shih, D.J. Katzmann, J.D. Schnell, M. Sutanto, S.D. Emr, L. Hicke, Epsins and Vps27p/Hrs contain ubiquitin-binding domains that function in receptor endocytosis, Nat Cell Biol, 4 (2002) 389–393.

[65] P.S. Bilodeau, J.L. Urbanowski, S.C. Winistorfer, R.C. Piper, The Vps27p Hse1p complex binds ubiquitin and mediates endosomal protein sorting, Nat Cell Biol, 4 (2002) 534–539.

[66] W. Mobius, Y. Ohno-Iwashita, E.G. van Donselaar, V.M. Oorschot, Y. Shimada, T. Fujimoto, H.F. Heijnen, H.J. Geuze, J.W. Slot, Immunoelectron microscopic localization of cholesterol using biotinylated and non-cytolytic perfringolysin O, J Histochem Cytochem, 50 (2002) 43–55.

